# The usual and unusual functions of thioredoxins in the metabolism and stress-response of sulfate-reducing bacteria

**DOI:** 10.1101/2024.09.27.615472

**Authors:** Erica L.-W. Majumder, Liyuan Hou, Fawn B. Whittle, Sharien Fitriasari, Valentine V. Trotter, Gareth P. Butland, Chris Petzold, Judy D. Wall

## Abstract

Thioredoxins are small, universal, disulfide isomerase proteins with required functions in oxidative stress response and RNA synthesis, among others. However, little is known about how anaerobic organisms maintain their intracellular redox balance or how thioredoxins may function differently under anaerobic metabolism. In this study, we investigated the roles of thioredoxins in sulfate-reducing microorganisms (SRMs). SRMs use sulfate as their primary electron acceptor in respiration to produce sulfide and are found in various environments including marine, freshwater sediments, guts and biofilms on ferrous metals where corrosion occurs. We found SRMs lack common redox maintenance molecules and macromolecules but have many and varied thioredoxins belonging to three types. Then, we probed their functions in the model SRM, *Desulfovibrio vulgaris* Hildenborough (DvH), by an *in vivo* disulfide bond capture proteomics experiment in both non-stressed and oxidatively stressed conditions. Our results demonstrated that thioredoxin 1 (Trx1) was essential in DvH and selectively responded to oxidative stress. Our data supported its role in RNA synthesis and energy transduction since Trx1 interacted with DsrC and QmoB. Thioredoxin 3 (Trx3), an atypical thioredoxin, was observed to have roles in sulfur transfer and dissimilatory sulfur metabolism. Next, DvH thioredoxin system protein encoding genes were deleted and single deletion mutant strains were tested for growth phenotypes under a variety of different electron donors, acceptors and toxic metal stresses. It is found that dissimilatory sulfate reduction improves resistance of DvH to metal stress. It appeared the sulfide provided certain protection to DvH from silver and uranium stress.

**Importance:** We put forth new functions for thioredoxins and a more robust understanding of sulfate reducing microorganisms physiology. Thioredoxin is of general interest because it has been widely studied for redox homeostasis or cancer therapies dealing with the excess of reactive oxygen species (ROS). Our results indicated that these proteins do have functions in stress response, even in microorganisms that generate large amounts of sulfide. We also identified interaction partners for an atypical thioredoxin, suggesting distinct roles from conserved thioredoxin. Mechanisms of metal stress response were found to be different than direct oxidative stress. Thioredoxin did not appear to be involved in uranium reduction electron transfer pathways, contradicting a hypothesis from the literature.

## 1. Introduction

Thioredoxin proteins are members of the redoxin family of proteins and are highly conserved across all kingdoms of life(1). They are primarily involved in oxidative stress response or redox homeostasis, often in conjunction with low-molecular weight thiol molecules. In addition, these dithiol proteins have essential functions such as being the electron donor to Ribonucleotide Reductase (RNR) or directing in protein folding(2). Thioredoxin systems are composed of two proteins; thioredoxin, a small ∼12 kDa disulfide isomerase protein and a dedicated reductase(2), thioredoxin reductase (TrxR), whose electron donor is the pyridine nucleotide, NAD(P)H. Thioredoxins act as a disulfide isomerase, making or breaking disulfide bonds on their numerous substrates. The family is characterized by the thioredoxin fold structure and the active site containing the two enzymatic cysteines spaced apart by two additional amino acids. Thioredoxin system proteins have been a popular drug target for dealing with diseases of redox imbalance(3), such as the overproduction of reactive oxygen species in the mitochondria of cancer cells. Moreover, viruses can discriminate between bacteria, virus, and human cells by sensing thioredoxins and the presence of selenium in thioredoxin reductases(4, 5). They have also been well-studied in plant systems(6), generally with the objective of combating diseases as well. Thioredoxins and their interaction partners have the subject of several recent reviews(2, 3, 7–21).

The ubiquity and importance of thioredoxins in so many diverse biological processes have prompted higher throughput study of their native substrate proteins. Techniques applied include yeast two-hybrid studies in the early 2000’s to emerging proteomic techniques and large scale-tagging and pull-down efforts(22–31). An *in vivo* interactome study in *E. coli* by disulfide bond capture between thioredoxin and its native substrates identified over 90 new partners for that thioredoxin(22). A similar study in *Chlamydomonas* probed the entire thiol-based redox control systems including thioredoxins and demonstrated how widespread the redoxome is throughout the cell(23). These studies investigate the mechanisms of oxidative stress response in molecular detail. The roles of thioredoxins in these model aerobic organisms now include: oxidative stress response, DNA and RNA synthesis, biosynthesis of many cofactors and vitamins, redox based transcription and translational control, sulfur assimilation, post-translational modifications and more(32). Thioredoxins are also studied by inhibition of their activities by treatment with cadmium or silver(33), which have a high binding affinity to the thiol bonds of the thioredoxin active site cysteines. Thioredoxin impacts on metabolism are best known through the disulfide isomerase activity of the protein, but it is also known to be a subunit of protein complexes, an electron donor or, in some cases, provide non-thiol functions(16).

The role of redoxin family proteins in anaerobic environments has not been well characterized, despite the fact that anaerobes are more sensitive to oxygen and subsequently oxidative stress. In this study, we examine the roles of thioredoxins of the model anaerobic Sulfate-Reducing Bacterium (SRB) *Desulfovibrio vulgaris* Hildenborough (DvH). SRBs are part of a larger class of Sulfate-Reducing Microorganisms (SRM) that have a unique mode of respiration, dissimilatory sulfate-reduction (DSR). Sulfate respiration uses eight electrons to reduce sulfate producing sulfide via adenosine 5’ phosphosulfate (APS) and sulfite in a series of enzymatic reactions(34). SRM therefore live in a reducing environment with high concentrations of the toxic molecule sulfide. Their metabolism and excretion of copious amounts of sulfide also account for the majority of the corrosion damage they cause on ferrous pipelines and ships and the souring of oil, costing millions of dollars annually(35). SRM excrete sulfide in the gut microbiome that can act as a signaling molecule between the microbes and the host epithelial cell immune system(36). The excess sulfide around SRM may allow non-standard mechanisms of redox maintenance and different roles for sulfur-dependent proteins like thioredoxins compared to non-sulfate-reducing organisms. Previous work on DvH thioredoxins has shown them to preserve the overall thioredoxin fold structure but demonstrated unique substrates and an unusual active site(37). The expression of DvH thioredoxin genes has also been shown to respond to oxidative stress by microarray analysis(38).

In our investigation of the function of DvH thioredoxins, we show that SRM generally lack glutathione (∼92% of the 50+ genomes examined) and other low-molecular weight thiol molecules used for intracellular redox balance. We hypothesize that excreted sulfide may influence the redox buffer in SRM. We found that SRM possess many thioredoxins with diverse active sites in their genomes. We therefore generated single deletion mutants of DvH thioredoxin family proteins and measured their growth phenotypes under several electron donor and acceptor combinations and under metal stress conditions. We also tested the uranium reduction capability of DvH compared with the thioredoxin mutant strains. Then, we over-expressed mutants of thioredoxins where the second active site cysteine was converted to an alanine that allowed trapping of the native substrate partners in stable intermolecular disulfide bonds. This was performed for both thioredoxins of DvH in oxidatively-stressed and non-stressed conditions. We then identified the native substrate proteins with proteomics. In that way with both targeted and untargeted approaches, we investigated the roles that the thioredoxins play in the metabolism and growth of DvH. Thioredoxin 1 (Trx1) was found to have essential functions and be the primary responder to oxidative stress. Thioredoxin 3 (Trx3) has an atypical active site(^37, 39^) and had unusual interaction partners for sulfur transfer. Overall, we demonstrate that thioredoxins in DvH have both the known functions seen in other biological organisms, but also unusual functions not previously observed. Thioredoxins are crucial to the functioning of SRM and their survival in their unique anaerobic environment.

## 2. Materials and Methods

### 2.1 Bacterial strains, growth conditions and reagents

Cultures of all *Desulfovibrio vulgaris* Hildenborough strains (Table S1) were grown from freezer stocks housed at University of Missouri first on rich MOYLS4 medium (*40*) and then on defined MO medium (41) at 34°C in anaerobic growth chamber (Coy Laboratory Products Inc., Grass Lake, MI) with 95% nitrogen, 5% hydrogen, and oxygen level below 1.5% (average 0.9%) as described previously (42). Strains used as controls were wild type DvH, mutant parental strain DvH JW710, and DvH CAT401079(43) from previous studies. All chemicals were purchased from Sigma Aldrich (St. Louis, MO) unless otherwise indicated.

#### 2.1.1 Production of thioredoxin overexpression strains

Strains for the overexpression of thioredoxin proteins were generated by first building four plasmids (pMO3418-3421) with the backbone derived from a *Desulfovibrio* native plasmid (pBG1)(44). Plasmids and primers (Tables S2&S3) were designed in aPE (a Plasmid Editor)(45). Thioredoxin genes (*trx3* and *trx1*) were introduced after a high copy number promoter (Table S2). The Strep-TEV-Flag tag sequence needed for the pull-down step of the interactome experiment was obtained from a previous study(43, 46) and added to the plasmid maps for Trx1 and Trx3. To introduce the modified active site thioredoxins for the *in vivo* disulfide bond capture conditions, the second cysteine residue in the active site is converted to an alanine. The C33A and C34A mutations into DVU1839 and DVU0378 were introduced via primers during PCR. Plasmids were assembled by Sequence Ligation Independent Cloning (SLIC), transformed and then purified, following the reported method in De Leon et al.(47) Wild type DvH genomic DNA was the source of the templates for the thioredoxin genes. After purification, correct plasmids were verified by PCR analysis and Sanger sequencing. Purified plasmids were electroporated into DvH JW710 cells grown to an OD_600_ value of 0.6 and then allowed to recover overnight in 1 mL MOYLS4 medium with 100 μg/mL spectinomycin. Cultures that were resistant to spectinomycin were scaled up as described below for use in the interactome experiment. Passages were minimized in order to preserve the plasmid.

#### 2.1.2 Production of marker replacement mutants of thioredoxin system genes in DvH

Plasmids (pMO3402, 3404, 3408, 3410 and 3416) for producing single gene deletions of thioredoxin system genes in DvH by marker replacement (MR) were also constructed following the same SLIC, purification, transformation and screening procedure as described earlier (Table S2)(47). Plasmid pUC from DvH JW9003 served as the backbone portion(40). Primers are listed in Table S3. Purified and sequence verified plasmids were electroporated in DvH JW710 cells grown to an OD_600_ value of 0.6. After recovery, mutants were plated for a phenotypic screen on separate MOYLS4 medium plates with either G418, spectinomycin, fluorouracil (5FU) or no selection. The resistance marker [*aph*(3’)-II] provides resistance to both kanamycin and G418, but G418 is more effective for selection. Colonies passing the screen were picked, grown up in 500 µL of MOYLS4 medium and then plated for single colony isolation. Selected colonies were grown up in 25 mL MOLS4 medium with antibiotics and gDNA were extracted from cell pellets. Marker replacement mutants were verified by PCR analysis and confirmed by Southern blot. This process produced DvH strains JW3402 (marker replacement of *trxR3*), JW3404 (marker replacement of *trx3*), and JW3408 (marker replacement of *trxR1*). DvH strain JW3416 had been produced in a previous study following a similar method.(43) Isolated strains baring the correct marker replacement mutations were then stored in glycerol at −80°C.

### 2.2 Thioredoxin protein data mining, dendrogram building and homology modeling

To find thioredoxins, glutathione and peroxiredoxins in diverse SRM, amino acid sequences (*trx1* DVU1839, *E. coli trx1* WP_072217281.1, *trx3* DVU0378, *E. coli gshB* PJI64279.1 glutathione synthetase, peroxiredoxin *ahpC* DVU2247, and *E. coli* peroxiredoxin-partial WP_085445176.1) were searched by BLASTP into the genomes of sequenced SRM from the list provided by R. Rabus et al. (48). Thioredoxins types were assigned manually by examination of the amino acid sequence for the active site residues and whether there was a zinc binding site present at the N-terminus or not. A total of 113 thioredoxin sequences were identified from 41 SRM genomes. The amino acid sequence of all thioredoxins were aligned by Clustal Omega(49) and then a dendrogram was built using the tree builder in MegaX with 500 bootstrapped replicates(50). Tree was color-coded by the manual classification of thioredoxin family types (i.e., type 1, 2 and 3; Figure S1). Homology models of Trx1 and Trx3 were built on the Phyre2 server by entering the amino acid sequence of two DvH thioredoxins(51). The top scored model was selected for each. The reference structure for Trx1 is *E. coli* Trx1 PDB code 3DXB-chain E (52) and Trx3 was the previously solved NMR structure of DvH 0378 PDB code 2L6C(53). Models were visualized in Chimera (54) and the active site cysteines were highlighted (Figure 1 B and C). Identity scores were computed in the alignment tool of BLASTP.

**Figure 1:**
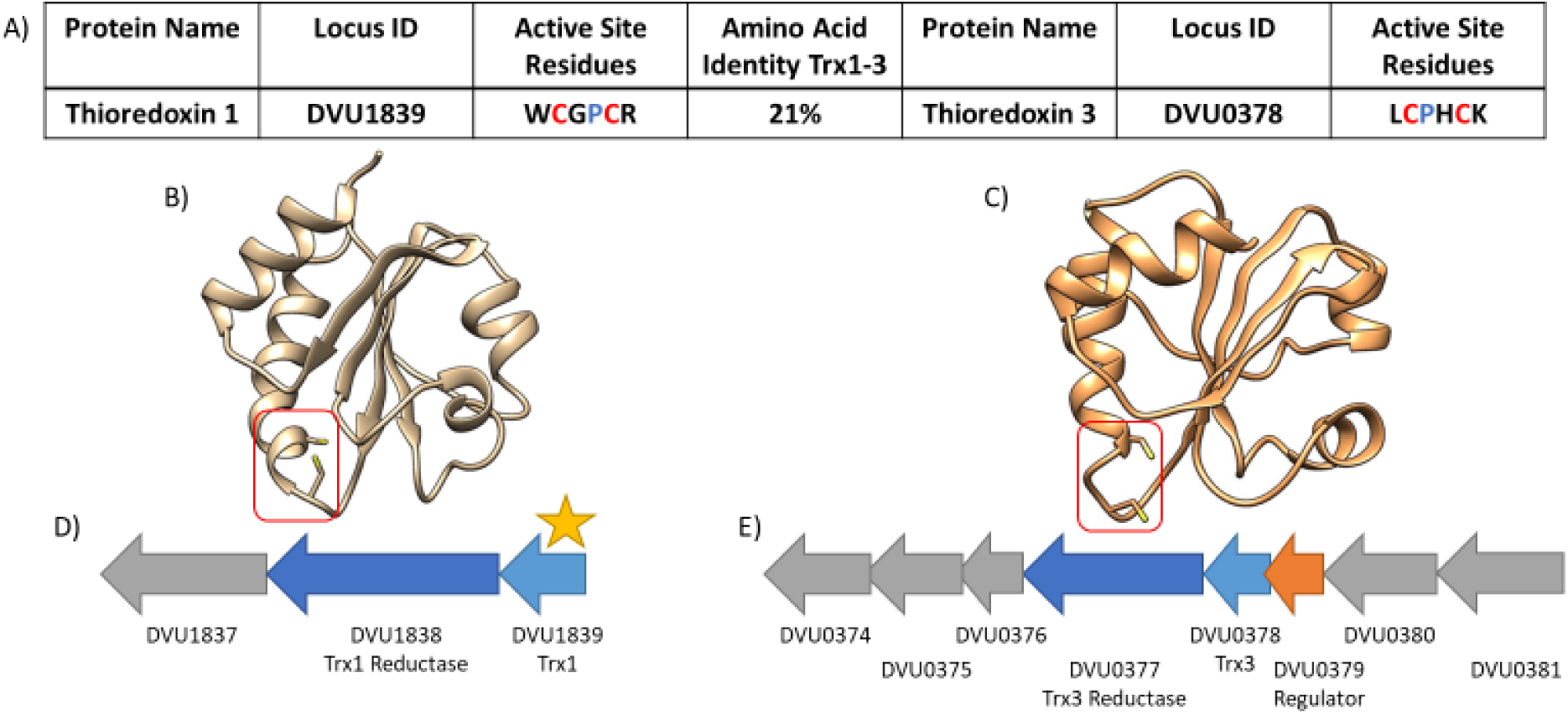
Thioredoxins of *Desulfovibrio vulgaris* Hildenborough. A) Amino acid content of active site and over all sequence identity between Trx1 and Trx3. B) Homology model of Trx1 on *E. coli* Trx1 (PDB code 3DXB-E). The active site is highlighted in red. C) Homology model of Trx3 on the NMR structure of DvH Trx3 (PDB code 2L6C). Operon structure of Trx1 (D) and Trx3 (E). Star indicates that a gene is predicted to be essential.

### 2.3 *In vivo* interactome of thioredoxins using overexpression strains

The DvH thioredoxin overexpression strains JW3418, JW3419, JW3420, and JW3421 were grown in triplicate in anaerobic conditions in MOYLS4 medium containing spectinomycin (Table S1). Wild Type DvH and DvH strain CAT401079, which has DVU1839 Trx1 STF-tagged on the chromosome, were also grown in the same way but without antibiotic as a control. Then, cultures were scaled up daily when OD_600_ reached 0.6 to 50 mL, 250 mL, then 1 L in MOLS4 medium. When the strains in 1L reached late-log phase (OD_600_=0.6-0.8), each replicate was divided in half into two separate bottles: one bottle for the non-stressed condition and one bottle for the oxidative stressed condition. To apply the oxidative stress, hydrogen peroxide to a final concentration of 1 mM was added to the respective bottles and mixed. All bottles were incubated anaerobically for another 2 hours. Bottles were then removed from the anaerobic chamber and cells were immediately harvested by centrifugation at 12,000 × g for 10 min at 4°C. The cell pellets were quickly frozen at −20°C for further analysis by mass spectrometry.

Cell pellets from the overexpression of thioredoxins were thawed and thioredoxins and their disulfide bonded interaction partners were purified as described in a previous work (43). Briefly, cells were resuspended in 5 mL lysis reagent BugBuster (Novagen, Darmstadt, Germany) per 1 g wet cells with the addition of 10 mM iodoacetamide (IAA) to prevent disulfide bond rearrangement. Cell debris were removed by centrifugation at 23,000 × g. Supernatant containing the solubilized proteins were incubated with Anti-Flag M2 magnetic beads for several hours at 4°C. Beads were loaded onto BioRad quick spin columns (BioRad, Hercules, Ca) and washed five times with AFC Buffer (10 mM Tris-Cl, 100 mM NaCl, 0.1% Triton X-100). Interaction partners were eluted by washing with Buffer A and 100 mM sodium dithionite to break the disulfide bonds. Eluant was collected and stored at 4°C. Columns were washed with Buffer A again and then TEV elution buffer was added containing TEV protease (50 mM Tris-Cl pH 7.9, 100 mM NaCl, 0.2 mM EDTA, 0.1% Triton X-100, 1 mM sodium dithionite and Complete protease inhibitor). Columns were incubated overnight at 4°C. Eluant was collected to verify whether desired thioredoxin was bound to the column or not. Proteins in the eluant were precipitated by a dose of acetone at −20°C and protein pellets were stored at −20°C until digestion and analysis by mass spectrometry.

Protein pellets were washed twice with 80% ice cold acetone and centrifuged at 12,000 × g for 10 minutes at 4°C between washes. The supernatant was removed and discarded. Protein pellets were dried for 5 minutes at room temperature. Then protein pellets were resuspended with 20 µL of 6 M urea, 2 M thiourea, and 100 mM ammonium bicarbonate. For quantitative comparisons, digestion volumes of all samples and replicates were normalized by protein abundance as measured with Pierce^TM^ BCA protein assay kit. For each sample, 1 µL of 200 mM dithiothreitol (DTT) in 100 mM ammonium bicarbonate was added and gently vortexed for 1 hr at room temperature to reduce disulfide bonds. The protein solution was then alkylated with 4 µL of 200 mM IAA in 100 mM ammonium bicarbonate, gently vortexed, wrapped up in aluminum foils, and incubated at room temperature for 1 hr. After alkylation an excess of 4 µL of 200 mM DTT in ammonium bicarbonate was added to neutralize the remaining IAA in solution with gentle vortexing for 1 hr. Urea concentration was reduced by adding 155 µL of Milli-Q water. This lowered the concentration of urea to approximately 0.6 M and thiourea to 0.2 M, allowing trypsin to be active in this concentration range. Trypsin solution (2 µg of trypsin in 20 µL 40mM ammonium bicarbonate with 10% acetonitrile (ACN)) was added to the sample. The sample was placed in a 37°C incubator for overnight digestion. After 16 hours, the sample was taken out and formic acid was added yielding 1% total concentration in the digested protein solution. Samples were then zip-tipped using Pierce^TM^ C18 Tips (100 µL; Prod # 87784) yielding sample in 35% ACN and 0.5% formic acid which was lyophilized.

21 uL 5% ACN, 1% formic acid in Milli-Q water was added to the lyophilized sample. Sample was vortexed and mixed by pipetting up and down and then centrifuged at 16,000 × g speed for 10 minutes and 21 uL of sample was transferred to an autosampler vial. An aliquot (18 uL) of each sample was loaded onto a C8 trap column (µ-Precolumn^TM^ – 300 µm i.d. × 5 mm, C8 PepMap 100, 5 µm, 100Å, ThermoFisher, Waltham, MA). Bound peptides were eluted from this trap column onto a 25 cm, 150 µm i.d. pulled-needle analytical column packed with HxSIL C18 reversed phase resin (Hamilton Co). Peptides were separated and eluted from the analytical column with a continuous gradient of CAN at 400 nL/min. The Proxeon Easy nLC HPLC is applied connected to an LTQ Orbitrap XL-ETD mass spectrometer. Mobile phase consisted of (A) 0.1% formic acid in water and (B) 99.9% ACN and 0.1% formic acid. The initial conditions of LC were 5% B and then was held for 2 minutes followed by 45 min ramp to 20% B. This is followed by a ramp from 20% B to 30% B in 50 minutes followed by a 15-minute rapid ramp to 90% B, followed by 90% B for 22 minutes. The gradient is then ramped down to 5% B for 1 minute and held at 5% B for 5 minutes. Total run time was 140 min. FTMS data were collected (30,000 resolution, 1 microscan, 300-1800m/z, profile) and then each cycle (approximately 3 seconds) the 9-most-abundant peptides (ignore +1 ions, >2000 counts) were selected for MSMS (2 m/z mass window, 35% normalized collision energy, centroid). Data was searched against the DvH proteome database (3385 sequences, update 1/17/2018) using sequestHT algorithm on Proteome Discoverer server. Matched peptides were uploaded into Scaffold and results were filtered by limiting the false discovery rate to 1%, increasing the minimum peptides to 4, using only exclusive unique spectral counts. Quantitative comparisons were done by grouping replicates and then applying a Fisher’s exact test with a cutoff of 0.05. Each mutated thioredoxin was compared to its not-mutated counterpart as the primary control. Comparison of Trx1 and Trx3, Trx1 under oxidative (Ox) stressed conditions vs. non-stressed condition (NS), and Trx3 under Ox stressed conditions vs. NS were carried out on the mutated samples. The WT and TEV elution controls were used to determine if a protein was a frequent flier and verify presence of correct bait protein. Passing proteins were exported to Excel and filtered for further removing frequent fliers, low averages, fewer than 2 replicates and proteins that only contained one cysteine residue in their sequence. The rest represented our high confidence list of thioredoxin interaction partners.

### 2.4 Single deletion mutant growth studies

2% inoculations of overnight cultures of each mutant with the single MR deletion for the strains JW3402, JW3404, JW3408, JW3416 and JW710 parental were grown anaerobically in capped, sealed, de-gassed with nitrogen and autoclaved Balch-type cultures tubes containing 5 mL of defined MO medium. All strains were grown in triplicate at 34°C. OD600 was measured at approximately 8 hours intervals. Additions to the base medium were at the concentrations as follows to make various combinations of electron donors and acceptors: 60 mM lactate, 60 mM pyruvate, 60 mM/1 mM formate/acetate, 30 mM sulfate, 20 mM sulfite, 20 mM thiosulfate, 2.5 mM cysteine, 2.5 mM sulfide, and 1 g/L yeast extract. Silver nitrate, cadmium chloride and uranyl acetate were all added to 1 mM.

### 2.5 Uranium reduction assay

Uranium reduction was monitored using an adaptation of the method by D.A. Elias et al.(55), (56). DvH cells from each strain tested (JW strains 3402, 3404, 3408, 3416 and 710 parental) were grown to late-log stage in triplicate on a defined medium in the Coy anaerobic chamber, as described in section 2.1. All reagents and consumables were incubated in the anaerobic chamber for at least before performing the assay. Cells were pelleted by centrifugation at 12,000 xg. Cells were washed in the same manner as the preparation for electroporation in section 2.2. Cells were then transferred back into the anaerobic chamber. They were then resuspended in equal amounts of anaerobic bicarbonate lactate buffer at pH 7, which was prepared the night before. Briefly, 2.5g of Sodium bicarbonate was mixed with 1L of sterilized deionized water and boiled under pressure and then transferred to the anaerobic chamber. The next day or when temperature had equilibrated, the pH was adjusted to a pH of 7 and sodium lactate was added to a final concentration of 10 mM. Cells for the heat-killed control were incubated in a dry bath at 80°C for 20 minutes. Each of the assays was started by adding uranyl acetate to a final concentration of 1 mM to tubes containing the resuspended cells, except the bicarbonate lactate buffer only control condition. Samples were taken for the assay of uranium reduction at 0, 1, 2, 4, 8, and 24 hours after addition of uranyl acetate. At each time point, 100 uL of each sample was collected and placed in a cuvette. Then 100 µL of complexing solution (25 g DCTA trans-1,2-Diaminocyclohexane-N,N,N,N-tetraacetic acid, 5 g sodium fluoride, 65 g sulfosalicylic acid in 1L of deionized water) was added immediately to quench the reaction. Then, 100 µL Tris buffer (149 g Tris base in 1L of water at pH 7.85), 500 µL ethanol 100 µL PADAP dye solution (0.05 g 2-(5-Bromo-2-pyridylazo)-5-(diethylamino)phenol in 100 ml reagent grade ethanol), 350 µL water, were added sequentially. Samples were incubated for 45 min at room temperature with light mixing and the absorbance was measured at 457 nm on a spectrophotometer. Absorbances were converted to uranium (VI) concentrations by a standard curve acquired with the same stock solution and assay buffer.

Data availability statement: All data are presented in the manuscript. Raw proteomics files and plasmid sequences are available upon request.

## 3. Results

### 3.1 Thioredoxin protein data mining, dendrogram building and homology modeling

To investigate the roles of thioredoxins in SRM, we first determined if thioredoxins were present in diverse SRM genomes by blasting known thioredoxin amino acid sequences and then recorded the type of thioredoxin and number found among the genomes. Thioredoxins of type 1 and 2 were found by blasting DvH *trx1* (DVU1839) in the SRM genomes and type 3 by blasting DvH *trx3* (DVU0378). Active sites were found by sequence alignment and manually inspected and categorized. We investigated 51 species from 12 bacterial families and one archaeal family that were identified as SRM in a study by Rabus and coworkers(57). All genomes contained at least one *trx* and 121 thioredoxin genes were detected in total based on sequence alignment yielding an average of 2.4 thioredoxin genes per species (Table 1). The maximum number of thioredoxin genes in an SRM genome was seven thioredoxin genes in *Desulfosporosinus orientis* DSM765T. In the detected thioredoxin sequences, we observed a variation in the amino acid residues surrounding the two primary cysteines and in extra metal binding sites in the N-terminus of the sequence. We therefore classified thioredoxins into three types based upon their active site structure and the presence or absence of the N-terminal zinc binding site. Similar classifications have been applied in other organisms, and our type 1 and type 2 classifications meet the following criteria. Thioredoxin type 1 (Trx1) family contains the classical thioredoxin protein fold and active site with the amino acid sequence WCGPCR, tryptophan-cysteine-glycine-proline-cysteine-arginine (or lysine). Trx1 in both protein fold and active site sequence is highly conserved across all kingdoms of life. Thioredoxin type 2 (Trx2) family contains thioredoxin type 1 as the C-terminal domain and has an additional zinc binding site near the N-terminus. In our classification, the thioredoxin type 3 (Trx3) family designation covers the rest of the thioredoxins that are not type 1 nor type 2. Type 3 therefore does not have the type 1 conserved active site or the Zn-binding site or N-terminal additional residues but does have the typical thioredoxin protein fold and an active site centered on 2 cysteines with two intermediate residues. However, the active site in thioredoxin type 3 can have many combinations of amino acid residues around the two cysteines. In total, 57 *trx1*, 14 *trx2* and 50 *trx3* were found. We generated a dendrogram of all found SPR thioredoxins by aligning the amino acid sequences in MEGA with muscle and found that the branching pattern agreed with our classification system (Figure S1). All SRM genomes probed contained a *trx1*, suggesting that this highly conserved protein is essential in SRMs. 40 of the 51 genomes contained two or more thioredoxins which indicates that a single thioredoxin gene may not be sufficient, that thioredoxins of type 2 and 3 may have different functions than type 1, and that a diversity of thioredoxin active sites is beneficial to the fitness of the microorganism.

**Table 1:** Thioredoxins of sulfate-reducing microorganism genomes.

| SRMs by Family | NCBI Taxonomy ID | GSH<br>-Syn | Peroxi-<br>redoxi<br>n | Trx<br>1 | Trx<br>2 | Trx<br>3 |
| --- | --- | --- | --- | --- | --- | --- |
| ***Desulfobacteraceae*** |  |  |  |  |  |  |
| *Desulfobacterium*<br>*autotrophicum* HRM2 | 177437 | - | + | 1 | 0 | 0 |
| *delta proteobacterium* NaphS2<br>(now unclassified) | 88274 | + | + | 2 | 0 | 1? |
| *Desulfatibacillum*<br>*alkenivorans* AK-01 | 439235 | - | + | 1 | 0 | 0 |
| *Desulfobacula toluolica* Tol2 | 651182 | - | + | 2 | 0 | 0 |
| *Desulfobacula* sp. TS | 1232437 | + | + | 1 | 1 | 1 |
| *Desulfotignum*<br>*phosphitoxidans* | 190898 | + | + | 1 | 1 | 0 |
| *Desulfococcus multivorans*<br>DSM 2059 | 1121405 | - | + | 1 | 1 | 0 |
| *Desulfococcus biacutus* | 125602 | - | - | 6? | 0 | 0 |
| ***Syntrophaceae*** |  |  |  |  |  |  |
| *Desulfobacca acetoxidans* | 60893 | - | + | 1 | 0 | 1 |
| ***Desulfoarculaceae*** |  |  |  |  |  |  |
| *Desulfarculus baarsii* 2st 14 | 644282 | - | - | 1 | 1 | 0 |
| ***Syntrophobacteraceae*** |  |  |  |  |  |  |
| *Syntrophobacter fumaroxidans*<br>MPOB | 335543 | - | + | 1 | 0 | 0 |
| ***Desulfobulbaceae*** |  |  |  |  |  |  |
| *Desulfotalea psychrophila*<br>LSv54 | 177439 | - | - | 1 | 0 | 1 |
| *Desulfobulbus propionicus* | 894 | - | + | 4 | 0 | 0 |
| *Desulfocapsa sulfexigens*<br>SB164P1 | 1167006 | ++ | + | 1 | 1 | 2* |
| ***Desulfovibrionaceae*** |  |  |  |  |  |  |
| *Desulfovibrio vulgaris* str.<br>Hildenborough | 882 | - | - | 1 | 0 | 1 |
| *Desulfovibrio magneticus* RS-1 | 573370 | - | + | 1 | 1 | 1 |
| *Desulfovibrio* sp. A2 | 298701 | - | + | 1 | 0 | 2 |
| *Desulfovibrio alaskensis* G20 | 207559 | - | - | 1 | 1 | 2 |
| *Desulfovibrio desulfuricans*<br>ND132 | 641491 | - | + | 1 | 0 | 0 |
| *Desulfovibrio africanus* str.<br>Walvis Bay | 690850 | - | + | 1 | 1 | 0 |
| *Desulfovibrio africanus* PCS | 1262666 | - | + | 1 | 1 | 2 |
| *Desulfovibrio alkalitolerans* | 293256 | - | + | 1 | 1 | 2 |
| *Desulfovibrio* sp. X2 | 941449 | - | + | 1 | 1 | 0 |
| *Desulfovibrio piezophilus*<br>C1TLV30 | 1322246 | - | + | 1 | 1 | 1 |
| *Desulfovibrio hydrothermalis*<br>AM13 | 1121451 | - | + | 1 | 0 | 0 |
| *Desulfovibrio gigas* ATCC<br>19364 | 1121448 | - | + | 4 | 0 | 0 |
| *Desulfovibrio aespoeensis*<br>Aspo-2 | 643562 | - | + | 1 | 1 | 1 |
| ***Desulfonatratonumaceae*** |  |  |  |  |  |  |
| *Desulfonatratonum*<br>*thiodismutans* | 159290 | - | + | 1 | 0 | 2 |
| ***Desulfohalobiaceae*** |  |  |  |  |  |  |
| *Desulfohalobium retbaense* | 45663 | - | + | 2 | 0 | 0 |
| ***Desulfomicrobiaceae*** |  |  |  |  |  |  |
| *Desulfomicrobium baculatum* | 899 | - | + | 1 | 1 | 1 |
| ***Thermodesulfobacteriaceae*** |  |  |  |  |  |  |
| *Thermodesulfatator indicus*<br>CIR29812 (now own phylum) | 667014 | - | + | 1 | 0 | 0 |
| *Thermodesulfobacterium*<br>*geofontis* OPF15 | 795359 | - | - | 1 | 0 | 0 |
| *Thermodesulfobacterium*<br>*commune* DSM 2178 | 289377 | - | - | 1 | 0 | 0 |
| ***Peptococcaceae (Gram-positive SRB)*** |  |  |  |  |  |  |
| *Desulfotomaculum*<br>*acetoxidans* | 58138 | - | + | 1 | 0 | 1 |
| *Desulfotomaculum reducens*<br>MI-1 | 349161 | - | - | 2 | 0 | 0 |
| *Desulfotomaculum ruminis* DL | 696281 | - | + | 2 | 0 | 0 |
| *Desulfotomaculum*<br>*hydrothermale* Lam5 = DSM<br>18033 | 1121428 | - | + | 1 | 0 | 1 |
| *Desulfotomaculum kuznetsovii*<br>17 | 760568 | - | - | 2 | 0 | 0 |
| *Desulfotomaculum gibsoniae* | 102134 | - | + | 1 | 0 | 0 |
| *Desulfotomaculum nigrificans*<br>DSM 574 | 696369 | - | - | 2 | 0 | 0 |
| *Desulfotomaculum*<br>*carboxydivorans* DSM 14880 | 868595 | - | - | 2 | 0 | 0 |
| *Desulfosporosinus* sp. OT | 913865 | - | - | 0 | 0 | 4 |
| *Desulfosporosinus orientis*<br>DSM 765 | 768706 | - | + | 0 | 0 | 7 |
| *Desulfosporosinus youngiae*<br>DSM 17734 | 768710 | - | + | 0 | 0 | 3 |
| *Desulfosporosinus meridiei* | 768704 | - |  | 0 | 0 | 3 |
| DSM 13257 |  |  |  |  |  |  |
| *Desulfosporosinus acidiphilus*<br>DSM 22704 | 646529 | - | + | 0 | 0 | 2 |
| *Candidatus Desulforudis*<br>*audaxviator* | 471827 | - | - | 1 | 0 | 0 |
| ***Nitrospiraceae*** |  |  |  |  |  |  |
| *Thermodesulfovibrio*<br>*yellowstonii* DSM 11347 | 289376 | - | + | 1 | 0 | 0 |
| ***Archaeoglobaceae*** |  |  |  |  |  |  |
| *Archaeoglobus fulgidus* VC-16 | 224325 | - | + | 0 | 0 | 3 |
| *Archaeoglobus profundus* | 84156 | - | + | 1 | 0 | 1 |
| *Archaeoglobus sulfaticallidus*<br>PM70-1 | 387631 | - | - | 0 | 0 | 4 |

In addition to thioredoxins, other members of the redoxin family proteins are typically present and essential in cells. We therefore searched the SRM genomes for the presence of peroxiredoxins and glutathione synthetase to study the presence and whether roles of redoxin family proteins are different in SRMs. Since the intracellular environment of SRMs is highly reducing and sulfidic as a byproduct of dissimilatory sulfate reduction, we hypothesized that the landscape of redoxin proteins and their metabolites, which rely on active sulfur residues for functionality, would be different in SRM. Peroxiredoxin (Prx), as a small thioredoxin-like protein that specifically reduces peroxide species, was found in the majority SRM genomes (36 of 51), which indicates that most SRM retain the ability to reduce peroxides. We also searched for glutathione synthetase, which is responsible for the production of glutathione (GSH). Our results found that SRM lacked glutathione synthetase, with just four exceptions (Table 1). We also cataloged the known low-molecular weight thiols from any organism in the literature and did not find evidence of their occurrence in SRM (Table S4). Therefore, although classical thioredoxins and peroxiredoxins are present, the abundance and diversity of thioredoxins and the lack of GSH point to a different system for redox homeostasis in SRM. Due to the absence of GSH or other LMWT’s, we hypothesize that thioredoxins or other members of the redoxin family are filling this role of redox balance.

To test this hypothesis, we investigated the thioredoxin systems of a model SRM, DvH, with genome mining and experiments. DvH has two thioredoxins annotated in the genome that are only 21% identical to each other via amino acid sequence (Figure 1A). Global gene expression studies of DvH showed that both *trx1* and *trx3* are highly expressed, and Trx1 was among the most abundant sequences. NMR structures have been solved previously (52, 53) and homology models were generated based on these NMR structures to visualize the active site conformation (Figure 1 B&C). Trx1 has the canonical thioredoxin protein fold and active site, WCGPCR (Figure 1A &B). Previous work demonstrated that it had normal disulfide isomerase and reductase activity, but also identified pyruvate: ferredoxin oxidoreductase (PFOR) as a substrate, which was a novel finding for any trx1. Trx1 begins a short operon containing its dedication thioredoxin reductase (TrxR1) and another protein of unknown function (Figure 1D). Trx1 is predicted to be essential from gene fitness assays conducted in other studies. The fitness value is 0, indicating it is not possible to make transposon mutant in the gene of *trx1*.(58) The other thioredoxin in the DvH genome is designated as Trx3 because it is neither a type 1 nor a type 2 thioredoxin. Trx3 has the normal fold, but has a very atypical active site where the proline residue has been relocated compared to typical thioredoxin sequences. The sulfur groups of the active site cysteines rotated so that they no longer pointed to each other. The divergence of Trx3 and Trx1 suggests a very different disulfide isomerase mechanism in Trx3 (Figure 1 A&C). In line with the observation of the structural active site, the previous work showed that Tx3 did not have normal thioredoxin enzyme activity and its function was not determined (37, 39). The operon structure of Trx3 also deviates greatly from Trx1 (Figure 1E). Trx3 is in the middle of an operon, followed by its dedicated reductase and the presumed regulator, which is cAMP-dependent. A Thioredoxin 3 homolog was also previously implicated in the uranium reduction electron transfer pathway of related SRM *Desulfovibrio alaskensis* G20(59). In the dendrogram of all mined SRM thioredoxin sequences, the overall branching pattern was driven by the sequences of the active site. DvH Trx3 grouped with other unusual trx3, but was closest to the *trx3* homolog of *D. alaskensis* G20 (Figure S1, arrows). DvH Trx1 was found among other highly similar and conserved *trx1* (Figure S1, arrows). Our protein alignments and modeling in conjunction with earlier studies suggest that thioredoxins are essential in DvH and may fill the void in the absence of other typical redox regulators. We furthermore hypothesize that Trx3 functions differently than Trx1 due to the difference in the active site.

### 3.2 *In vivo* interactome

We investigated the native interaction partners of both Trx1 and Trx3 from DvH with an *in vivo* proteomics assay. Substrate proteins were captured in a stable intermolecular disulfide bond on thioredoxins during growth by introducing a stable overexpression plasmid with a thioredoxin where the second cysteine of the active site was mutated to alanine, preventing the full isomerase reaction. The wild type DvH thioredoxins were left intact on the chromosome because of the essentiality of Trx1. By placing the mutated strains on a high copy number plasmid, we would outcompete wild type (WT) expression levels. Due to their role in oxidative stress response, both Trx1 and Trx3 were tested in non-stressed and oxidatively stressed conditions. After purification and selective elution of disulfide bonded proteins, substrate partners were digested with trypsin and their peptides measured by mass spectrometry. Peptides were then mapped to the DvH proteome. Trx1 native interaction partners were found by doing statistical comparisons between the mutated *trx1* and the not-mutated strain expressed on a plasmid (Table 2). Frequent fliers or non-specific binding proteins were identified from the WT and TEV elution controls. All identified proteins passed the filtering as outlined in the methods section.

**Table 2:**
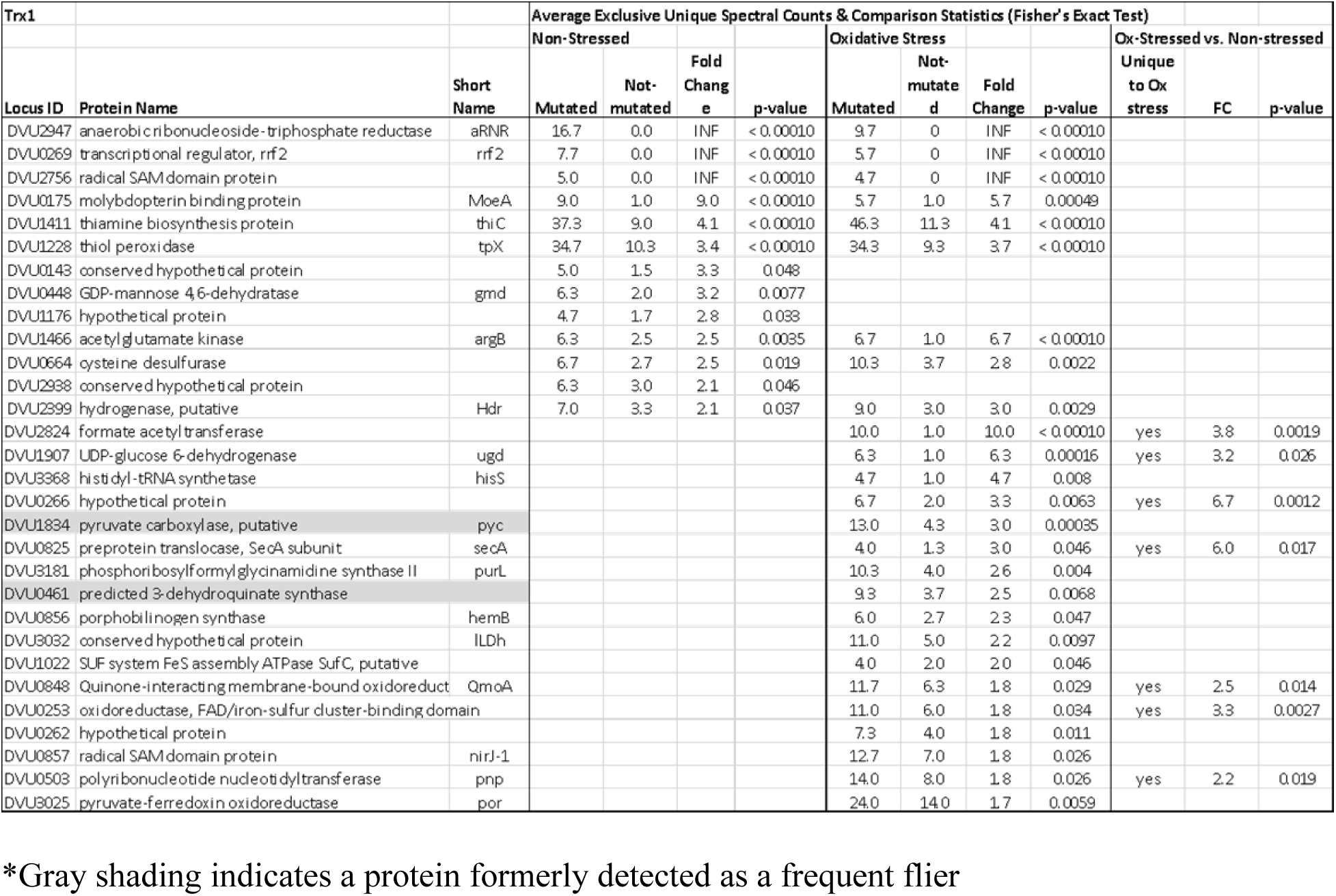
Interaction partners of DvH Trx1 (DVU1839) under non-stressed and oxidative stress conditions.

The results of the interactome study showed some trends. Under non-stressed conditions, DvH thioredoxin type 1 has around 10 partners that are identified from stringent filtering (Table 2 and Figure 2a). When considering partners seen in the other Trx1 controls, we find Trx1 has additional partners and especially those that are known in the literature (22, 60, 61). This is likely because the not-mutated thioredoxin overexpression and WT controls can still catch real interactions. This increases the number to almost 20 proteins, including DsrC and its dedicated reductase (Table S5). Interaction partners play roles in carbon metabolism, sulfur metabolism, amino acid biosynthesis, protein folding and so on (Figure 2A). Under oxidative stress conditions, the number of interaction partners doubled (Table 2), and many of the interaction partners under non-stressed conditions had the similar expression levels as those with known functions in oxidative stress response. Trx1 was only found with TrxR, and likewise Trx3 had a strong preference for TrxR3. TrxRi (DVU1457, thioredoxin reductase) was not detected in any data set in this study. Trx1 and Trx3 were detected in the controls, confirming the correct bait protein in each condition and the selectivity of the assay. Trx3 (Table 3 and Table S6) has far fewer interaction partners than Trx1 and did not show a significant response to oxidative stress. Trx3 partners revolve around protein translation and amino acid biosynthesis. When including the partners determined from WT controls, the range of partners is larger, which play roles in carbon metabolism and DNA synthesis. However, they did not show a response to oxidative stress. To our knowledge, this is the first report of substrate proteins for a thioredoxin with an atypical active site. To complete our analysis, we quantitatively compared the interactomes of Trx1 and Trx3 to determine if the interaction partners were unique or shared between the proteins Trx1 and Trx3 (Tables 4, S7 and Figure 2b). A few interaction partner proteins were shared between the two thioredoxins, but most were unique to one thioredoxin. Among the stringently filtered proteins, only one (i.e., DVU0462 chorismate mutase) showed preference of Trx3 over Trx1. Observed again by this analysis, Trx3 did not have a response to oxidative stress. We therefore will explore the interaction partners of DvH thioredoxins by functional class as follows.

**Figure 2:**
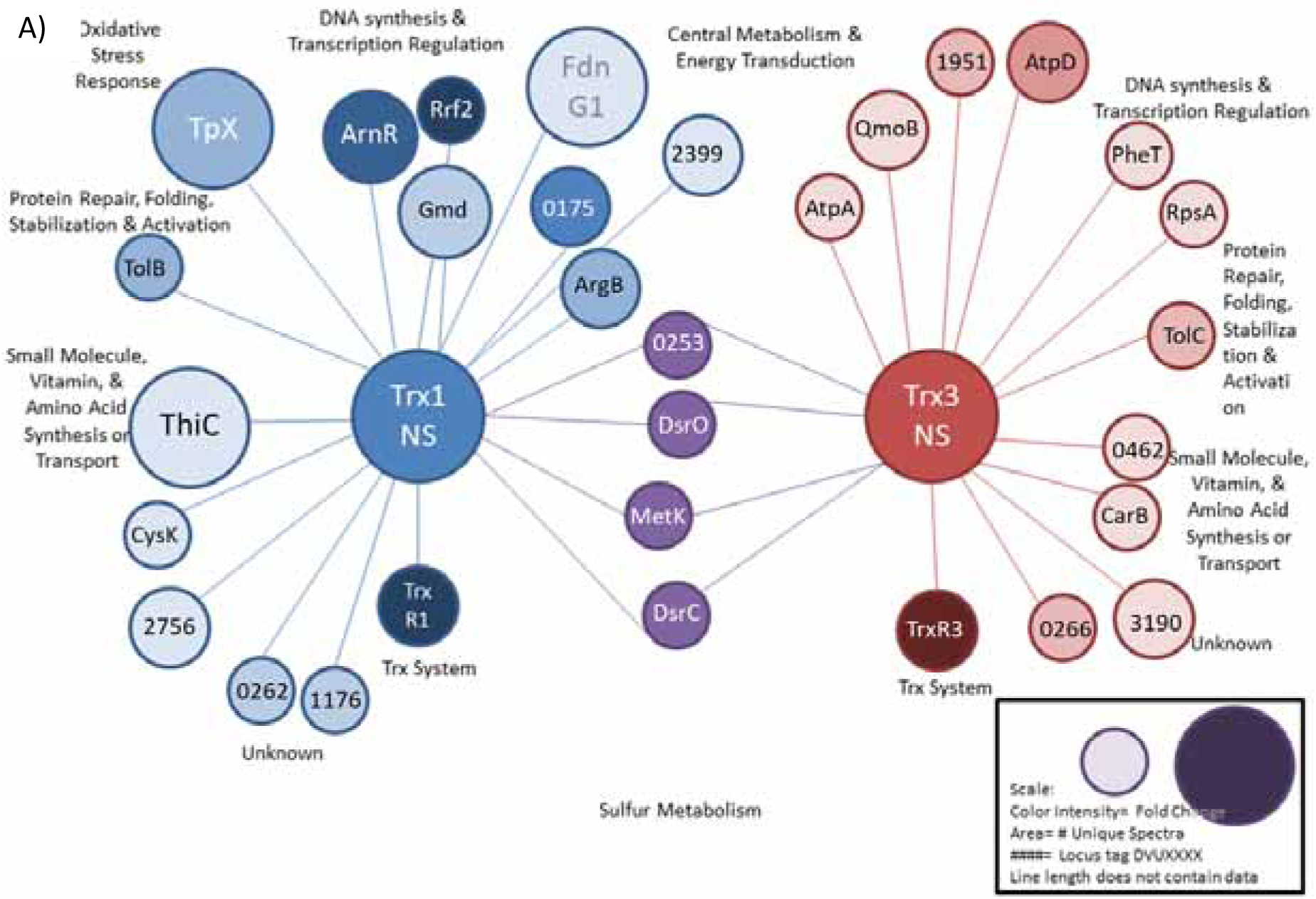
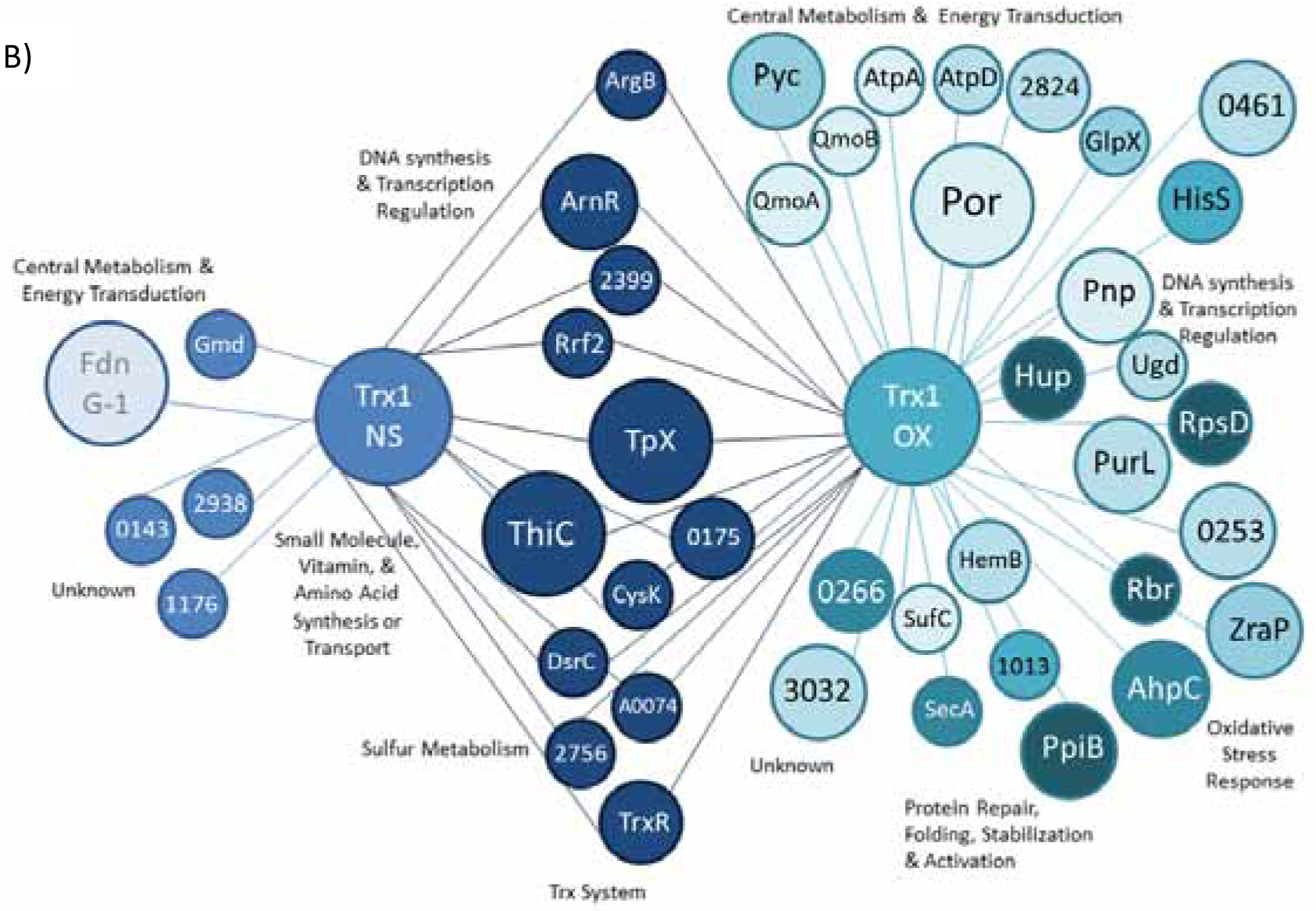
Visualization of thioredoxin interactome results. A) Trx1 interactome under non-stressed condition (blue) compared to Trx3 interactome under oxidative stressed condition (turquoise), as well as shared interaction partners (dark blue). B) Trx1 interactome under non-stressed condition (blue) compared to Trx3 under non-stressed condition (red), and shared interaction partners (purple). For both panels, a darker color represents a higher fold change with lower p-values. A larger size denotes more unique spectral counts per protein. Line length does not contain information.

**Table 3:** Interaction partners of Trx3 (DVU 0378) under non-stressed and oxidative stress conditions.

| Trx3 |  |  | Average Exclusive Unique Spectral Counts & Comparison Statistics (Fisher's Exact Test) |  |  |  |  |  |  |  |  |  |  |
| --- | --- | --- | --- | --- | --- | --- | --- | --- | --- | --- | --- | --- | --- |
|  |  |  | Non-Stressed |  |  |  | Oxidative Stress |  |  |  | Ox-stressed vs. Non-stressed |  |  |
| Locus ID | Protein Name | Short Name | Mutated | Not-mutated | Fold Change | p-value | Mutated | Not-mutated | Fold Change | p-value | unique to ns or ox | Fold Change | p-value |
| DVU2543 | hybrid cluster protein | hcp2 | 10.3 | 1.0 | 10.3 | < 0.00010 |  |  |  |  | NS | 5.2 | 0.00068 |
| DVU0732 | valyl-tRNA synthetase | valS | 5.0 | 1.0 | 5.0 | 0.0093 |  |  |  |  | NS | 10.0 | 0.0019 |
| DVU1025 | uracil phosphoribosyltransferase | upp | 5.0 | 1.7 | 3.0 | 0.045 |  |  |  |  |  |  |  |
| DVU0953 | tyrosyl-tRNA synthetase | tyrS | 7.0 | 2.7 | 2.6 | 0.032 |  |  |  |  |  |  |  |
| DVU0587 | formate dehydrogenase, alpha subunit | fdnG-1 | 10.0 | 4.0 | 2.5 | 0.015 |  |  |  |  |  |  |  |
| DVU3190 | hypothetical protein |  | 10.7 | 4.7 | 2.3 | 0.022 |  |  |  |  | NS | 2.5 | 0.045 |
| DVU1043 | GMP synthase | guaA | 10.0 | 4.7 | 2.1 | 0.038 |  |  |  |  |  |  |  |
| DVU3319 | proline dehydrogenase | putA | 14.3 | 7.7 | 1.9 | 0.041 |  |  |  |  |  |  |  |
| DVU1067 | membrane protein, Bmp family |  |  |  |  |  | 5.7 | 2.0 | 2.8 | 0.003 | OX | 2.4 | 0.0044 |
| DVU0462 | chorismate mutase |  |  |  |  |  | 5.0 | 2.7 | 1.9 | 0.029 |  |  |  |
| DVU2055 | methionyl-tRNA synthetase | metG |  |  |  |  | 5.3 | 3.0 | 1.8 | 0.031 |  |  |  |

**Table 4:** Significiant interaction partners of Trx1 compared to Trx3 under non-stressed and oxidative stress conditions.

| Trx1 vs Trx3 |  |  | Non-Stressed |  | Oxidative Stress |  |
| --- | --- | --- | --- | --- | --- | --- |
| Locus ID | Protein Name | Short Name | Fold Change | p-value | Fold Change | p-value |
| DVU2947 | anaerobic ribonucleoside-triphosphate reductase | aRNR | 8.3 | < 0.00010 | INF | < 0.00010 |
| DVU0269 | transcriptional regulator | rrf2 | INF | < 0.00010 | INF | 0.00016 |
| DVU2756 | radical SAM domain protein |  | 3.33 | 0.0018 | INF | 0.00074 |
| DVU0175 | W-formylmethanofuran dehydrogenase |  | 6.75 | < 0.00010 | 5.667 | 0.0012 |
| DVU1411 | thiamine biosynthesis protein | thiC | 4 | < 0.00010 | 5.56 | < 0.00010 |
| DVU1228 | thiol peroxidase | tpX | 5.777 | < 0.00010 | 6.867 | < 0.00010 |
| DVU0448 | GDP-mannose 4,6-dehydratase | gmd | 2.375 | 0.012 |  |  |
| DVU1176 | hypothetical protein |  | 4.67 | 0.0031 |  |  |
| DVU1466 | acetylglutamate kinase | argB | 3.167 | 0.0005 | 2.667 | 0.028 |
| DVU0664 | cysteine desulfurase |  | 4 | 0.00078 | 4.133 | 0.00059 |
| DVU2399 | hydrogenase, putative |  | 2.333 | 0.0094 | 5.4 | 0.0026 |
| DVU2824 | formate acetyltransferase |  |  |  | INF | < 0.00010 |
| DVU0825 | preprotein translocase, SecA | secA |  |  | 4 | 0.038 |
| DVU3032 | conserved hypothetical protein |  |  |  | 2.2 | 0.015 |
| DVU0462 | chorismate mutase |  | 0.266667 | 0.018 |  |  |

\*Blue shading denotes proteins that had higher interactions with Trx3 than with Trx1

#### A. Trx1&3 interaction partners involved in thiol mediated redox regulation (oxidation stress response, PTM’s, transcription factors)

Thioredoxins are commonly associated with oxidative stress response as their primary role in a cell. Our interactome experiment results are consistent with the notion that thioredoxins are required for redox stress. In the case of SRM, redox stress could be oxidative or reductive and the reversible nature of disulfide isomerase proteins would be able to handle that in either situation. Trx3 did not have a measurable response to oxidative stress as only three additional proteins were found in this condition, but the number of spectra was barely above the significance cut offline (Table 3). Likewise, none of Trx3’s partner proteins have known roles in oxidative stress response. The absence of oxidative stress response in Trx3 is attributed to the difference in active site. It seems that more conventional active sites are required for oxidative stress response and the non-canonical active site sequences may fill other roles in DvH cells.

Trx1, on the other hand, displayed the expected response to oxidative stress. The number of interaction partners increased and the number of times that interaction was detected also increased in stressed conditions (Table 2). Several of the highest binding interacting proteins during non-stress conditions had decreased spectral counts under oxidative stress, suggesting that Trx1 has different priorities in times of stress and suppresses other functions. Known stress response proteins which interacted with Trx1 include: Tpx (DVU1228 or thiol peroxidase/peroxiredoxin), AhpC, zraP, rbO, rbr2. Most of these proteins are in the redoxin family and have a characteristic thioredoxin fold and can be reduced by thioredoxin reductases or thioredoxins. Thioredoxin is known to reduce Tpx(62) but others had not as yet been reported to interact with Trx1. Beyond these stress response proteins, thioredoxins also function in stress response by controlling transcriptional regulators and post-translationally modifying proteins based on the redox balance and thiol content of the cell. This function of thioredoxins was also observed in our interactome results. One of the most observed interactions of Trx1 was with a poorly characterized regulator from the rrf2 family, TmcR. In DvH, DVU0269 is named TmcR because it was found to be a part of and regulated the operon producing the tmc protein complex, which is involved in energy transduction(63). Rrf2 contains active site cysteine residues that in some members of the family are part of a thiol/disulfide switch(64–66). Oxidizing agents form disulfide bonds on rrf2 can increase thioredoxins expression in some cases(67). DVU0529 is also annotated as an rrf2 which was characterized by Voordouw and coworkers(68). Trx1 was also associated with DNA binding protein, hup2, which was found only under oxidative stress conditions, given further suggestions of thiol and thioredoxin mediated control of transcription in DvH, especially in response to oxidative stress.

In addition to making and breaking disulfide bonds, thioredoxins have been recently associated with thiol-based post translational modifications (PTMs) of proteins, driving disulfide bond formation, S-desulfhydration and S-nitrosylation. Many other PTMs are sulfur mediated like others such as S-lipidation and are likely to have a thioredoxin component. In our studies, we observed DVU1834 as an interaction partner of Trx1. DVU1834 is annotated as pyruvate carboxylase (pyc), which has been noted as a frequent flier. However, in our study, pyc was above the controls for frequent fliers, and thus suggests it is a real interaction. Ju and coworkers in 2016 also observed a direct interaction of Trx and Pyc and thioredoxin was involved in protein desulfhydration(69). Trx3 has an interaction partner involved in the PTM S-nitrosylation. Hybrid cluster protein (Hcp2, DVU2543) has been documented in other organisms in response to nitric oxide (NO) or reactive nitrogen species (RNS) stress(70). The interaction with Trx and Hcp2 has a direct role in the PTM and protection of the cell against RNS(71). The presence of these PTMs are further indicators of the major role sulfur plays in regulating a myriad of functions in the DvH cell. There are also sulfotransferases and methyltransferases associated with thioredoxins that in addition to assembling amino acids and cofactors could also mediate some of the sulfur based PTMs. Aligned with such thioredoxome studies (22, 60, 61), the range of proteins and genes that are subject to redox control or direct interactions with thioredoxins is very large.

#### B. Trx1&3 interaction partners involved in biosynthesis of DNA, amino acids, cofactors and vitamins and protein translation and translocation

The majority of proteins associated with thioredoxins in DvH are involved in the biosynthesis of many aspects of the cell from DNA, RNA, amino acids, proteins, EPS sugars, cofactors and metal clusters (Figure 2). The extent of thioredoxin functions and their contacts underlie many essential functions throughout the cell. Protein folding, assembly, electron donation, and sulfur source transport are all carried out in some ways by thioredoxins.

In addition to the fact that thioredoxins have a hand in regulating transcription, they are involved in the biosynthesis of DNA and RNA precursors, such as purine and pyrimidine nucleotides. A well-documented of thioredoxins in many organisms is to serve as the electron donor/subunit for ribonucleotide reductase (RNR) and therefore thioredoxins are involved in the regulation of the total rate of DNA synthesis in the cell especially during the cell division and DNA repair since RNR is the rate-limiting enzyme in this process(8). Trx1 was found with the triphosphate form of anaerobic RNR (aRNR) DVU2947, but not significantly with the diphosphate aRNR DVU3379. This may be because this is an essential function of thioredoxins and the WT chromosomal Trx1 may have a preferential bound. Regardless, this is strong evidence that the thioredoxin fulfills this essential role in DvH. Under oxidative stress, Trx1 was also found with PurL from the purine biosynthesis pathway, which has a documented interaction with thioredoxin in other organisms and links to methanogenesis(72). Trx3 interacted with upp (DVU1025), which is not only involved in pyrimidine metabolism but also used as a genetic marker tool(42, 73). Trx3 was also bound to guaA (DUV1043), which plays roles in purine biosynthesis with other biological molecules. Thioredoxins are heavily involved in the synthesis of nucleotides and their polymer chains.

Many of the documented interaction partners of Trx1 and Trx3 were involved in the biosynthesis of amino acids. Cysteine and methionine biosynthesis are common and well-documented roles for thioredoxins in many organisms. However, one thing where SRM are quite different from their non-dissimilatory sulfate reducing counterparts is that their biosynthesis of cysteine and methionine is different(74). Despite that SRM like DvH use different enzymes, we still see an association of thioredoxins with these enzymes, suggesting that thioredoxins still have a role to play in biosynthesis of cysteine and methionine. The primary evidence is the interaction of Trx1 with hypothetical protein DVU2938, which was recently found to be a missing member of the new methionine biosynthesis pathway in DvH(74). Trx is known to carry out similar reactions to the chemistry that this enzyme is suggested to convert aspartate semialdehyde to homocysteine. ArgB (DVU1466), an enzyme in amino acid biosynthesis of arginine and others, is also a partner of Trx1. Interestingly, a homolog to this protein was found to interact with the thioredoxin in yeast mitochondria, but in a manner that was not dependent on the active site cysteine thiol residues(75). Other trx1 interaction partners associated with amino acid biosynthesis are hypothetical protein DVU0143 and the apparently essential DVU0461 or 3-dihydroquinate synthase on the tryptophan biosynthesis pathway. DVU0461 appears to be an essential gene(74) and the biosynthesis pathway DVU0461 belongs to is subject to redox regulation(6). Trx3 was found associated with several tRNA synthetases and putA (DVU3319) on the proline biosynthesis and metabolism pathway, which has been previously confirmed to interact with thioredoxins and promote stress survival(76). DVU0462, along with other members of the shikimate biosynthesis pathway that is discussed below, was another specific partner of trx3 associated with amino acid biosynthesis. Thioredoxin is known to mediate this protein and pathway in other systems.

Beyond the building blocks of proteins, thioredoxins were also found to have associations with proteins involved in translation, protein folding and protein translocation. Both Trx1 and Trx3 were bound to the tRNA synthetase enzymes of several amino acids, but this interaction was much more common for Trx3. DVU3368, hisS, which is also involved in tRNA and protein translation(77) was found with Trx1 primarily under oxidative stress conditions. Similar results were observed in the *E. coli*(22). As is documented in *E. coli*, ribosomal subunits were detected as interaction partners(22). DVU0503, pnp, is involved in mRNA degradation and was previously seen as a Trx interaction partner in cyanobacteria. This indicates this protein can be functioned as transcription factors under the thiol-based redox control(78). Disulfide bond formation is a key step in protein folding and assembly for many proteins. It is likely that some of the interactions we observed in our study were related to protein folding and assembly, but it is difficult to decipher each protein’s function. In other organisms, thioredoxin has associations with chaperone proteins. We also observed chaperone proteins in our study. However, these interactions did not pass our stringent filtering as they have been treated as frequent fliers previously. So, it is possible that chaperone interactions with thioredoxins exist, despite being at approximately equal amounts in all conditions we tested. We did find direct evidence of thioredoxin mediating protein translocation with at least two interaction partners, SecA with Trx1 and ABC-transporter of lipoproteins with Trx3, under oxidative stress conditions. SecA belongs to a complex involved in protein translocases in the cytoplasmic membrane as the peripheral motor subunit, where thioredoxin may be acting as the electron donor or block(79, 80) or moved by the complex(81).

Thioredoxin also has a major role in the assembly of many protein cofactors, especially the cofactors containing a metal. Thioredoxin was found to be imperative for the assembly of iron-sulfur clusters and tetrapyrroles including molybdopterin. Thioredoxin has been observed to mediate Fe-S formation and DvH is no exception(82). Interaction partners included NifS (Dvu0064) which brings the sulfur atom to some assemblies(83), SufC (DVU1022) which is an interaction partner of TrxQ in *Staph aureus* (*73*) and shown for Fe-S biogenesis(84), and hcp2 (DVU2543) which was with the prismane FeS cluster in nitrite reduction and RNS. The presence and quantity of Fe-S associated interaction partners increased under oxidative stress. As for the heme and tetrapyrrole biosynthesis, two members of this operon were found with *hemB* (DVU0856) and *nirJ-1* (DVU0857). NirJ-1 has been observed to require an unusual heme biosynthesis route in methanogens(85). HemB DVU0856, like other members of this pathway have been found associated with thioredoxins and thiol-based redox control has been demonstrated to be of the utmost importance for this pathway(86–88). In addition to making the heme precursor, these proteins were associated with response to RNS stresses(89). MoeA (DVU0175), molybdopterin biosynthesis enzyme, did not have a lot of homology to other things with other SRMs, but the homolog ddes_1824 responded to NO stress(89). GuaA (DVU1043) is also associated with cobalamin biosynthesis.

The amino acid biosynthesis pathways often link into metabolic pathways associated with the biosynthesis of other necessary metabolites like vitamins and non-metallic cofactors that were also heavily associated with thioredoxins. Many of these proteins are radical SAM enzymes. Many of other sulfur-containing cofactors and their metabolites were dependent on thioredoxins: Thiamine (ThiC DVU1411), quinones (ThiH-like DVU2756), folic acid (DVU3032), and shikimate pathway (DVU0461, DVU0462-*trx3*). ThiC carries out radical SAM chemistry with diverse S-donors(90), and thioredoxins could be the sulfur shuttle. Many thioredoxin partners observed carry out radical SAM chemistry indicating an important role for thioredoxins in sulfur and methyl transfer reactions. Thioredoxin has been implicated in the production of quinones(91). For instance, a deletion mutant of TrxB1 in *Lactobacillus* upregulated menaquinone biosynthesis(92). Also, a homology of DVU2756 was found associated with the thioredoxin in *E. coli*(*25*). It’s worth noting that DVU0461 went with Trx1 and the next protein DVU0462 on the operon went with Trx3 very selectively, suggesting interesting shikimate pathway interactions.

Another documented interaction of thioredoxins is with sugar metabolism. Unlike in *E. coli* where interactions were more involved in glycolysis and gluconeogenesis, the DvH thioredoxin-sugar metabolism interactions centered on the production of sugars for extracellular polymeric substances (EPS) or external signaling molecules. These trx1 interaction partners were gmd (DVU0448) and ugd (DVU1907), which thioredoxins may reduce the active cysteines. Gmd in DvH was upregulated during phase transition and carbon flux(93). A homolog to udg in plants has an interaction with a thioredoxin that coordinated the Calvin cycle and the pentose phosphate pathway(94).

#### C. Trx1&3 Interaction partners involved in energy transduction, carbon, electron donor metabolism

Energy generation from carbon metabolism, sulfur metabolism and energy transduction were also found as thioredoxin associations. Thioredoxins already had known interactions with pyruvate and lactate complexes in DvH and other organisms. Thioredoxin is also the sulfur carrier for 3’-Phosphoadenosine-5’-phosphosulfate (PAPS) reductase in assimilatory sulfate reduction and mediates thiosulfate reduction or oxidation when thiosulfate is used as an alternative sulfur source in dissimilatory or assimilatory sulfate reduction. In DvH, thioredoxin was found with transmembrane complexes involved in energy transduction via electron transport. It was found that the Tmc operon was a dominate interaction as well as two subunits of Qmo and Hdr-operon member DVU2399. Tmc operon members with a thioredoxin interaction were: DVU0262, DVU0266 and DVU0269. DVU0269 is the tmcR regulator that has been discussed above. DVU0266 and DVU0262 were found under oxidative stress. DVU0266 is a hypothetical protein with 6-bladed beta propeller, tolB like domain, and homolog DvMF_2557 had a strong negative gene fitness phenotype under starvation. DVU0262 is also a hypothetical protein but belongs to a two-component response regulator protein family whose homologs are associated with synthrophic growth and phosphate limitation survival in *D. desulfuricans*(*95, 96*). DVU2399 has an electron transfer domain and is cytoplasmic so thioredoxin may be involved in reducing the iron-cofactors and is highly upregulated under synthrophic growth in DvH(97) and *D. alaskanesis* (98). This indicates the role of thioredoxins in energy generation and the metabolic flexibility of SRM. Trx1 was found with QmoA and QmoB (DVU0848 and DVU0849), a heterodisulfide reductase(41). The nature of interaction with thioredoxins is unknown, but the links with these proteins further suggest the role of thioredoxins in energy transduction and the link with sulfur metabolism.

Carbon and electron donor metabolisms of lactate, pyruvate and formate proteins were also observed as interaction partners of Trx1 and Trx3. Lactate metabolism thioredoxin interaction partners are DVU3032 that has homologs acting as L-lactate dehydrogenase(99) and DVU0253 involving in atypical lactate utilization(100). DVU3032 has strong fitness phenotypes under no vitamins and lactate as a carbon source and DVU3032 seems to be important for lactate utilization(101). DVU0253 has a ferredoxin-like domain, and therefore the thioredoxin interaction with this protein would likely be redox-based(93, 99, 100). Pyruvate metabolism was seen with pyruvate ferredoxin oxidoreductase (por, DVU3025), pyc (DVU1834) and pyruvate-formate lyase (DVU2824). Por or PFOR was previously detected as an interaction partner of Trx1 in DvH(37). Pyc was downregulated in high pH in DvH(102) and characterized as a frequent flier in a global DvH proteome pull down study(43). DVU2824 and its homologs in other SRMs have been strongly associated with fermentative growth(103, 104), again demonstrating the metabolic flexibility of these SRMs and the key roles of thioredoxins contributing to that ability(105, 106). Thioredoxin has been shown to act as an electron donor to the Fe-S cluster of DVU2824 homologs in other organisms(107). Interestingly, RNR nucleotide reduction is coupled to oxidation of formate to carbon dioxide, where thioredoxin also serves as the electron donor(108). Thus, thioredoxin could be the electron donor mediating both processes-supplying energy on both sides to drive reaction. Formate utilization was observed with Trx3 partner fdnG-1 (DVU0587). FdnG-1 is a cytoplasmic selenocysteine-containing protein associated with formate utilization and metabolic flexibility(109). TrxR is required to put selenium into fdn, and as such finding a thioredoxin interaction is not unexpected with this protein(110).

Dissimilatory sulfate reduction enzymes such as the Dsr subunits DsrC and DsrO were also observed as interaction partners of thioredoxin. This indicated a newly observed function of Trx, as Dsr enzymes are unique to SRM. It has been proposed that DsrC carries the sulfur from sulfate as a trisulfide to the membrane portion of the Dsr complex, bringing electrons and therefore linking DSR to ATP production(111–115). However, how the electron or sulfur is transferred from the trisulfide is unknown. This interaction suggests that thioredoxin might mediate the sulfur or electron transfer or in regenerating the subunit to receive the trisulfide. Trx is also known in thiosulfate metabolism. In DvH, the evidence of the interaction of Trx1 with DVU0143, whose homologs are SQR and conserved in Sox bacteria for thiosulfate oxidation(116). Many of the interaction partners in this section play roles as alternative sources of energy and metabolism employed by SRM such as syntrophy, methanogenesis, diverse nitrogen sources. This corresponds with the idea that the diversity of thioredoxins is related to the metabolic flexibility of this diverse microbial family.

### 3.3 Single deletion mutant growth studies

To further study the role of thioredoxins in the metabolism and stress response of DvH, single deletion marker replacement mutant strains of DvH were made by removing individual genes (i.e., *trxR1*, *trxR3*, *trx3* and the regulator of the *trx3* operon). Deletion of *trx1* was attempted but failed, likely due to the essentiality of *trx1* to DvH under the growth conditions tested in this study. To determine the role of thioredoxins in anaerobic respiration, specifically for carbon and sulfur metabolism, DvH was grown on several different electron acceptor and donor combinations (Figure 3). The ratio of donor to acceptor was held constant so that all conditions would have the same amount of potential energy for respiration and growth. The Trx3 operon mutants displayed a significant lag phase under lactate sulfate conditions, which were the standard donor acceptor combination used for growing SRB in bench-scale experiments (Figure 3A). Interestingly, deletion of TrxR1 did not impact growth, despite being the only known reductase for the putatively essential Trx1 (Figure 3). The lag phase in the *Δtrx3* mutants was observed again with growth on thiosulfate (Figure 3C), but the phenotype was rescued by growth on sulfite (Figure 3B), suggesting that Trx3 plays a non-essential but important role in dissimilatory sulfate reduction upstream of sulfite reduction. Trx1 is the known carrier of PAPS to PAPS reductase in assimilatory sulfate reduction (ASR) pathways (117, 118), but the Trx3 mediating this type of sulfur transfer process in the early sulfate reduction steps has not been observed. In the interactome experiment, Trx3 was found to interact with other sulfur transfer proteins, suggesting it may be a carrier of APS in dissimilatory sulfate reduction (Figure 2A, Tables S6). Thioredoxins have also been predicted in a pathway for thiosulfate reduction as an assisting protein to the thiosulfate reductase enzyme, further suggesting that Trx3 play roles in sulfate reduction (116, 119, 120).

**Figure 3:**
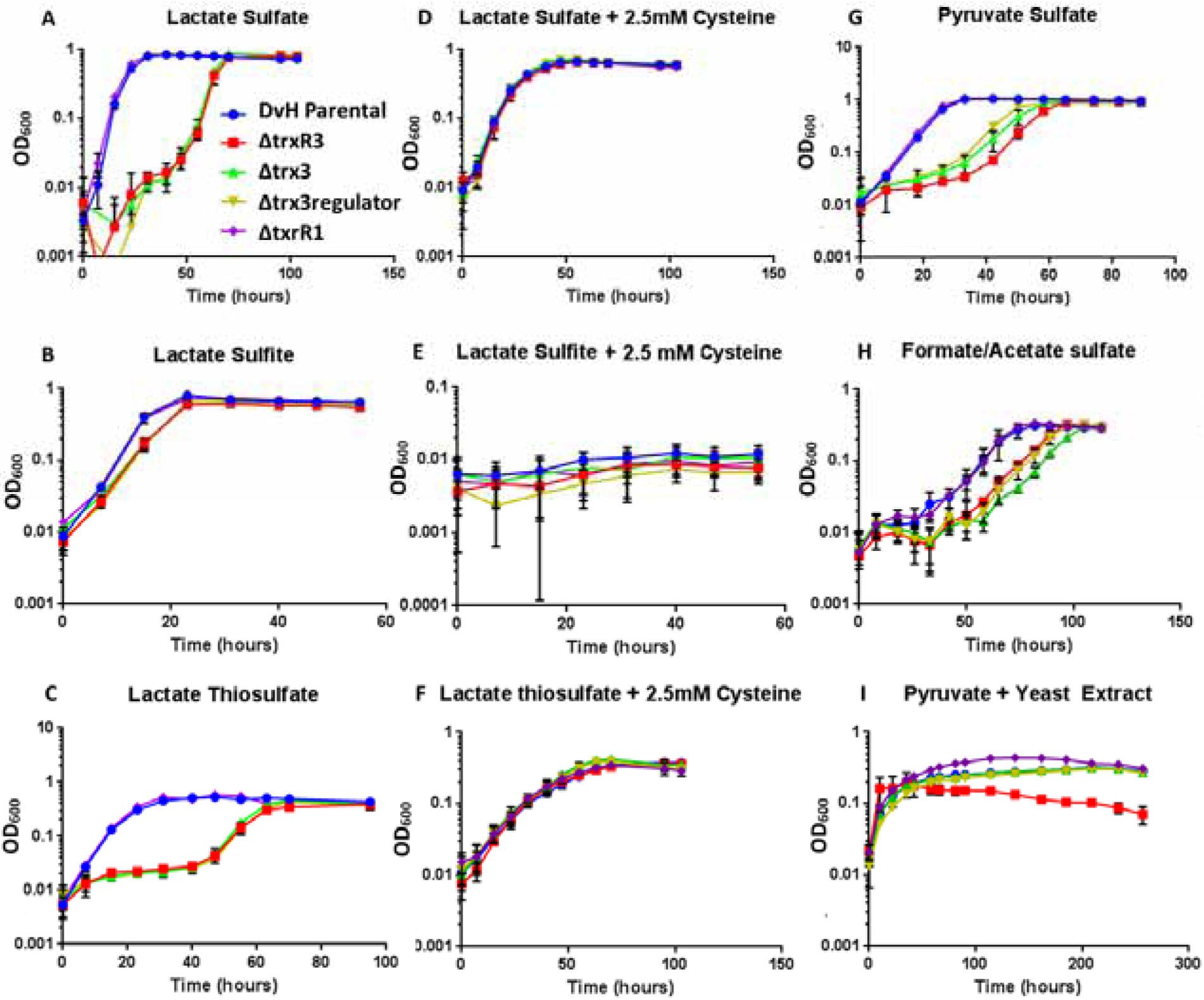
Growth behavior by OD_600_ of different single deletion mutants on various combinations of electron donors and acceptors. A-H) were performing dissimilatory sulfate reduction and I) was growing by pyruvate fermentation.

Investigation of Trx3 in dissimilatory or ASR was tested with cysteine addition to the same growth conditions in the previous paragraph (Figure 3 D-F). The cysteine amendments provided an additional sulfur source that bypassed DSR and provided the end product of ASR, bypassing that pathway. Therefore, Trx3 operon lag may not exist with cysteine amendment. Consistent with this idea, the addition of cysteine to the growth medium rescued the *Δtrx3* operon lag for lactate sulfate and thiosulfate (Figure 3 D and F). The addition of cysteine to the sulfite cultures resulted in no growth, which may be due to radical chemistry (Figure 3E). To determine if the rescued *Δtrx3* lag phenotype was from additional reducing power or a function of cysteine as a sulfur or amino acid source, the equimolar sulfide was added without cysteine in an additional growth experiment (Figure S3). The cultures with sulfide added displayed the ΔTrx3 operon lag phenotype suggesting that the cysteine rescue was not from additional reducing equivalents, but from bypassing the ASR and DSR processes before sulfite reduction.

Since Trx1 had been previously characterized to use PFOR as a substrate and in other organisms Trx1 is known to serve as an essential subunit of PFOR(37), growth on pyruvate as the electron donor and with sulfate as the electron acceptor as we have tested (Figure 3G). *ΔtrxR1* mutant did not display a growth phenotype which was contrary to our predictions, but the *Δtrx3* operon mutant had the lag phase. We also assayed if the role of Trx1 with PFOR would be present under fermentation rather than DSR respiration. Growth by pyruvate fermentation in DvH was initially more rapid, but plateaus quicker at a much lower maximum OD than DSR growth (Figure 3I). Pyruvate fermentation did not display a lag phase by any of these mutants, but *ΔtrxR3* did not reach the same maximum OD as the other mutants (Figure 3I). DvH can also use formate as an electron donor. Growth on formate/acetate sulfate was slower and reached a lower OD than other electron donors (Figure 3H). The *Δtrx3* operon lag phenotype was less pronounced, but still significant. Deletion growth studies on different electron acceptor and donor combinations suggested that thioredoxins, particularly Trx3 have a role in carbon and sulfur metabolisms. However, Trx1 functions could not be proved by this method.

In addition to their roles in carbon and sulfur metabolisms, thioredoxins are also known to be primary actors in responding to oxidative stress, as it was found in the results of the interactome experiments for Trx1. A source of oxidative stress that SRM experience in the environment is metals. Therefore, we tested the growth of the thioredoxin deletion mutant strains under different metal stresses (Figure 4). Cadmium and silver were chosen because they have been used in the literature as selective thioredoxin inhibitors in other organisms (33, 121). Despite this, SRM are known to make silver nanoparticles. Uranium was chosen because of the report that Trx3 is essential for uranium reduction(59). Also, SRMs are common members of field communities in uranium contaminated sites. We cultured the thioredoxin mutants under metal stress in both dissimilatory sulfate reducing conditions (lactate/sulfate) and fermentation conditions (pyruvate only). Fermentation conditions were employed because the sulfide produced by DSR has a strong affinity to bind metals and may therefore protect the cells from metal stress externally. This could mask the intracellular stress response with the involvement of thioredoxins.

**Figure 4:**
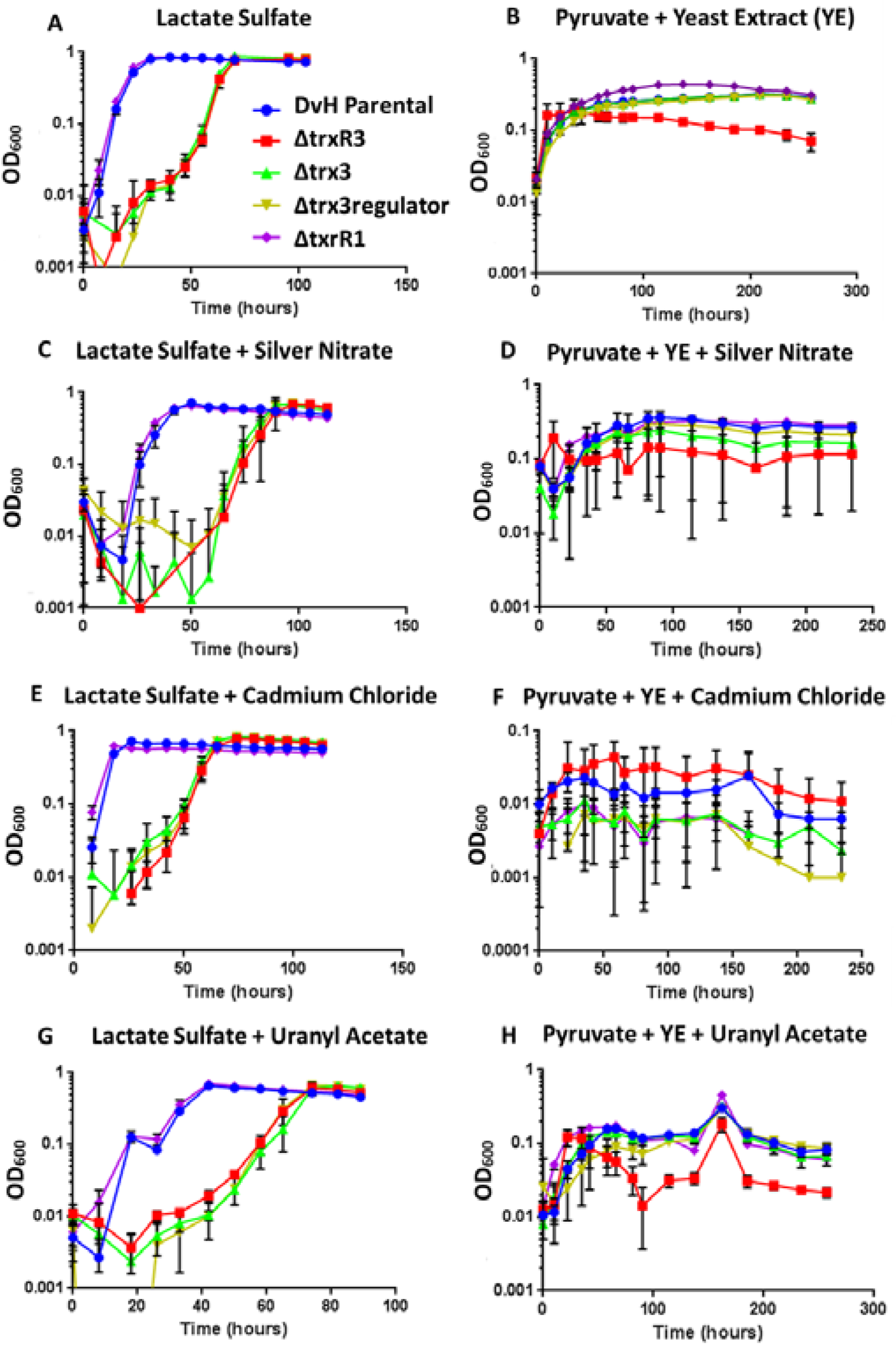
Single Deletion Mutant Growth Curves Under Metal Stress. Growth behavior monitored at OD_600_ of different single deletion mutants under various metal stresses in triplicate. All metal additions are at 1 mM. Panels A,C,E&G are growing by dissimilatory sulfate reduction while panels B,D,F and H are growing by pyruvate fermentation.

Under sulfate reducing conditions with the concentration of metals tested, the metals did not exhibit visible stress on the cells or alter the growth phenotypes (Figures 3, 4C, 4E, and 4G). *Δtrx3* operon mutants still displayed a growth lag with metals present comparing to no metal stress conditions. Growth lagged slightly in the presence of silver (Figure 4C, while the presence of cadmium gave a slight increase in the initial growth rate (Figure 4E). Growth in the presence of uranyl acetate was also slowed more than with silver and cadmium (Figure 4G). Although uranium respiration was tested, DvH did not grow on uranium as the sole electron acceptor (Figure 4G inset). Single thioredoxin deletion mutants were also tested for their ability to reduce uranium as Trx3 has previously been shown to be involved. The reduction of U(VI) to U(IV) was monitored. In our study, all mutants displayed the same capacity as parental DvH to reduce U(VI) (Figure S2). Under sulfate-reducing conditions, the parental DvH strain was not inhibited by either thioredoxin inhibitors, cadmium or silver (Figure 4 C&E). Since Trx1 is essential in DvH, the inhibition of Trx1 by the metals should result in a non-growth phenotype in all strains including the parental strains. This lack of thioredoxin inhibition by known inhibitory metals may result from the metals which are first bounded to the excreted sulfide and do not enter the cytoplasm.

To test the hypothesis that the excreted sulfide from DSR is protecting the cells from metal stress, the same strains were grown under pyruvate fermentation conditions in the presence of the same concentrations of metals (Figure 4 B, D, F, and H). In the absence of excess sulfide, the pyruvate fermentation conditions displayed different responses to the metals with different mutants. The growth with silver and uranium was relatively similar to the non-metal treated samples, but did display a growth lag phase and eventually reaching approximately the similar OD values (Figure 4 D&H). It appears the sulfide was lending some protection to DvH cells from silver and uranium stress, where the cells were able to recover. The *ΔtrxR3* mutant struggled the most, especially under uranium stress, in a manner distinct from the non-stress condition. The effects of sulfide on the uranium reduction assay was eliminated by washing cells with sulfate free buffer before the assay (Figure S2). Growth on cadmium was severely impaired in all strains including the parental strain (Figure 4F). This observation is in line with our prediction of growth inhibition by a thioredoxin inhibitor molecule. Then, in the absence of sulfide, cadmium may be acting as a Trx1 inhibitor in DvH. This further suggests that excreted sulfide acts as a protectant against oxidative and metal stresses for SRMs enabling their documented ability to live in metal-contaminated sites and in the oxic/anoxic transition zone of sediments.

## 4. Discussion

### 4.1 SRMs lack common redox maintenance molecules and macromolecules

For most organisms including mammals, the main method of maintaining redox homeostasis is to keep a high intracellular concentration of the LWMT GSH. In bacterial species that lack GSH, other LWMTs have been identified (Table S4). However, based on our genome-based searching and previous chemical studies, neither GSH nor other LMWTs have been found in SRMs. Our analysis revealed that SRM lack the genes for glutathione synthetase, and only four of the over 50 genomes had the gene (Table 1). In agreement with our genome-based evidence, previous studies looking for GSH or a replacement molecule in SRM did not detect GSH or another low-molecular weight thiol (LMWT) that can act as an alternative to GSH (37, 53, 122). The trend held for archaeon and gram-positive bacterial phyla along with the more classical SRMs, delta-proteobacteria (Table 1). This suggests that SRMs use a different method to maintain redox balance in their cells. This may not be that surprising given that SRMs are unique in that they live in a sea of free thiol, i.e. sulfide, where concentrations can be in the milimolar range both intra and extracellularly. By comparison, aerobically grown *E. coli* maintains GSH at 10 mM. Therefore, since free sulfide can act as a reductant in the same way as other LMWTs at similar concentrations, the free thiol alone may act as the redox buffer in SRMs. SRMs have a novel mechanism of redox homeostasis that is yet to be fully characterized.

### 4.2 Dissimilatory sulfate reduction improves resistance to metal stress

Another interesting observation from our batch experiments of growth studies was that the resistance of DvH mutants to metal stress was dependent on the mode of anaerobic respiration. Since there was no significant change with the metals on the behavior of individual mutants, it suggests the impact of metal stress is from the presence or absence of excreted sulfide. Cationic transition metals are known to have a high binding affinity with thiols and sulfide. Therefore, we propose that sulfide is binding the toxic metals either extra or intracellularly and preventing the metal from inhibiting thioredoxin or exerting its toxic effects. It also suggests that the literature-asserted specific thioredoxin inhibitors are specific for thiols or disulfide rather than thioredoxin alone. Cadmium in particular was far more toxic under pyruvate conditions, which may be due to the inhibition of trx1. These results further suggest that for some bacteria, metals like silver may not be an effective antibiotic. We also saw fewer interactions of thioredoxins with metal-associated proteins in DvH than in *E. coli* suggesting that sulfide may act on metals to enhance the resistance of metal stress in other SRMs. SRMs could also be exploited to make novel metal-sulfide nanoparticles. DSR makes SRM more robust in extreme environments.

### 4.3 SRPs possess diverse thioredoxins which are universal and required

The presence of multiple thioredoxins in all genomes queried suggest that thioredoxins are universal and required for organismal survival (Human tx1 and DvH Trx1 are 35% identical, Figure S4). Based on our thioredoxin interactome studies and thioredoxin interactome studies of other organisms, we observed that there are multiple roles of thioredoxins in biology. The number of thioredoxins per genome and active site diversity points to having multiple and different substrate proteins that are essential for cell survival across SRMs. This amount of diversity was surprising because to date, only plants are discussed as having many and varied thioredoxins in their complex genomes. The branching pattern of an amino acid sequence dendrogram confirmed our classification of the thioredoxin proteins into three types based on sequence variation in the active site, in agreement with the convention used in other organisms (11, 12, 123, 124). We should note that discovery of thioredoxins from sequence blasting may be less accurate in the genomes that were not closed. This can be exemplified by *Desulfococcus biacutus* which was not found with any thioredoxins in the unclosed reported genome, but it has six potential thioredoxin sequences when re-analyzed after using more recent genome assembly programs to close the genome. Our exploration of DvH Trx1 and Trx3 strongly suggests that differences in active site structure yield different functions for the protein in the cell’s metabolism. Diverse thioredoxins are also associated with metabolic flexibility in our study.

However, given the excess free thiol intracellularly, it begs the question as to the need for thioredoxin and other redoxin family proteins and their quantity *in vivo* in SRMs: Why is the selective thiol-mediated chemistry of redoxin proteins necessary in the presence of excess radical thiols? Sulfide produced at the end of DSR respiration is S^2-^. However, HS^-^ can act as a thiol radical sulfide depending on the pH. This thiol radical should readily bind with any free cysteine or methionine residues, and other radicals such as reactive oxygen species (ROS) and RNS. Metals could break disulfide bonds based on its reduction potential. Since it is a small diffusive and highly reactive molecule, sulfide could bind available sites much more quickly than proteins like thioredoxins. Yet, our studies in DvH demonstrate that Trx1 is essential and has many functions beyond redox maintenance. For instance, Trx1 could act as a subunit of aRNR, which can be replaced by sulfide. Similarly, Trx3 is acting as a sulfur carrier for DSR enzymes in a way that free thiol could not play a similar role because the binding of APS to sulfide would not be reversible and therefore could not be transferred to the receiving enzyme. The fact the disulfide isomerase activity can be carried out in the presence of excess thiol means that SRMs have a novel method to protect or sequester or repair efficiently these essential active site cysteines from bombardment by the thiol radical sulfide. The mechanism of disulfide chemistry in the presence of excess thiol, in conjunction with the mechanism of sulfate import and sulfide excretion which are also unknown in SRM, could be the subject of future investigations.

### 4.4 Trx1 has essential roles in transcription, translation, cofactor biosynthesis and oxidative stress response

Unlike other organisms, Trx1 in DvH is putatively essential. In our work with DvH, attempts at deleting or interrupting the trx1 gene were lethal, implicating that the gene is essential for survival under the conditions we applied. Thioredoxin systems in other organisms may be reduced by glutaredoxins, GSH or other related redoxin family proteins largely absent in SRM. For instance, Trx1 is not essential in *E. coli*, and the essential jobs of Trx1 would be filled by another protein. In DvH, deletion or mutation of Trx1 is likely not rescued because the genome lacks another Trx with the same active site. Therefore, Trx1 is required to carry the essential functions of thioredoxin in DvH. Trx1 has a well-documented essential function as the electron donor to RNR in aerobic organisms including mammals. This function was confirmed in DvH by the significant association between Trx1 and aRNR in the *in vivo* interactome results. Trx1 was also found with transcriptional regulator rrf2 and other nucleotide, amino acid, cofactors, and vitamins biosynthesis proteins confirming an important role for Trx1 in transcription and translation of DNA to RNA to proteins. Thioredoxins also mediated the protein folding and translocation of enzymes as well as some classes of PTMs in DvH.

As predicted from the canonical role of Trx1 family proteins, Trx1 had a strong response to oxidative stress induced by hydrogen peroxide. Trx1 had many more significant interaction partners under oxidative stress including AhpC and Tpx, which are common ones under oxidative stress. Thioredoxins also were associated with many proteins that respond to various nitrogen-species stress and metal stresses. SRMs are quite sensitive to inorganic nitrogen molecules, and this protective function of thioredoxins is likely important to their survival. The association with PTMs is also an important function of thioredoxins in oxidative stress response. Trx3 did not respond to oxidative stress, providing further evidence to the idea that different active sites result in segregation of duties. This assumption is further confirmed by the fact that Trx1 and Trx3 had very few interaction partners in common.

Trx1 was found only with the reductase on its operon, TrxR1, which is supported by previous enzyme activity assays(37, 39, 43). Although TrxRi (DVU1457) was not detected in any sample, the deletion of TrxR1 was able to grow at the same rate as the parental strain in all growth conditions suggesting that under loss of the preferred reductase conditions, TrxRi can substitute for TrxR1. This demonstrates functional redundancy within DvH redoxin family proteins to maintain the essential functions of Trx1 in the absence of its primary reductase. Trx1 had many associations with carbon metabolism and energy transduction complexes. Unfortunately, since Trx1 could not be deleted and TrxR1 was rescued in all growth conditions. Insight into the function of Trx1 in carbon metabolism and DSR respiration was not obtained from the growth studies of mutants. Despite this, we were able to investigate Trx1 involvement with literature evidence and the interactome results. A previous study using Trx1 as bait found that PFOR (DVU3025) was associated with DvH Trx1 (125). PFOR was also characterized as a ‘frequent flier’ in the global DvH pull-down assays, but it has been seen significantly in our targeted *in vivo* study. It should be noted that the DvH genome contains two different pyruvate ferredoxin oxidoreductases protein complexes. We observed the smaller atypical PFOR. This could be because of the position of the Trx1 as a subunit of PFOR that would prevent the affinity tag on Trx1 from binding to the column. Also, since the chromosome copy of trx1 was left intact in the genome, non-tagged and not-mutated Trx1 may have preferentially bound these essential partners, preventing us from observing them with our capture system. The close interactions between energy transducing complexes like Tmc(63), Qmo, Hdr and thioredoxins indicated that thioredoxins in DvH are involved in energy transduction and sulfur metabolism.

### 4.5 Trx3 has roles in sulfate reduction and protein translation

Trx3 had fewer but distinct interaction partners compared to trx1, which was expected from the atypical active site and unusual disulfide isomerase activity reported previously. Trx3 interacts with several tRNA synthetase proteins and SAM domain containing proteins suggesting its roles in protein translation and sulfur transfer. Trx3 did not have a measurable response to oxidative stress. Trx3 operon mutant strains displayed a lagging phenotype in many of the growth studies. Curiously, this phenotype could be rescued by the addition of cysteine, but not sulfide nor reductant. This suggested Trx3 was involved in providing organic or carbon bonded sulfur to DvH. Trx1 is the known carrier of PAPS during assimilatory sulfate reduction in mammals. The rescue on sulfite and addition of cysteine suggests that Trx3 may be the APS carrier and that DvH diverts some sulfide from DSR to ASR to produce cysteine. With an abundance of organic sulfur, the necessity of Trx3 may decrease. Furthermore, thioredoxins can chaperone thiosulfate in an alternative thiosulfate reduction step. This could explain the lag phase on thiosulfate or that the thiosulfate is being disproportionated into sulfate, which therefore had the same phenotype as the lactate sulfate condition. We attributed the lack of growth on the supplement with sulfite and cysteine to toxic molecules that may have formed from the reaction of sulfite and cysteine. The role of Trx3 in DSR is further supported by the absence of the lag phenotype under pyruvate fermentation conditions.

As mentioned above, the participation of Trx1 in DSR could not be probed by deletion mutant strains in DvH and, instead, Trx1 was overexpressed in the interactome experiment, leaving the native Trx1 intact. In our interactome results, Trx1, like Trx3, was also associated with a number of sulfur transfer proteins, including the trisulfide carrier DsrC. DsrC is proposed to be the electron carrier that links DSR to transmembrane potential for ATP synthesis via the Qmo enzyme and the trisulfide it brings, but the reductase of DsrC is not known. A different pull-down study found that DsrC was the only significant hit for Trx1 (43). In our data sets of Trx1 interaction partners, we observed both DsrC and QmoB, suggesting that Trx1 played a role in the energetic coupling of anaerobic respiration by reducing the trisulfide and freeing the electrons for use by Qmo. However, further studies are required to decipher if this association is a coincidence of available thiols from cysteine active sites or is serving the stated function in DvH.

### 4.6 Novel roles of thioredoxins in SRMs

Despite the reducing and sulfidic environment of the SRM cytoplasm, to our surprise, the overview picture of the global functions of thioredoxins in DvH highly resembles the functional associations found for thioredoxins in many other organisms. We did see many links to syntrophy, methanogenesis and nitrogen metabolism that are more restricted to microbes, which were more similar to SRMs. Also, thioredoxins may have the important ability to handle ROS and RNS. The most novel roles for thioredoxins in DvH were found to be related to the unique to SRM proteins for DSR and alternative biosynthesis of methionine. Therefore, we propose this revision to the SRM metabolic model in Figure 5. Thioredoxin was found to have multiple functions in almost every system in the metabolism, growth and defense of life for DvH.

**Figure 5:**
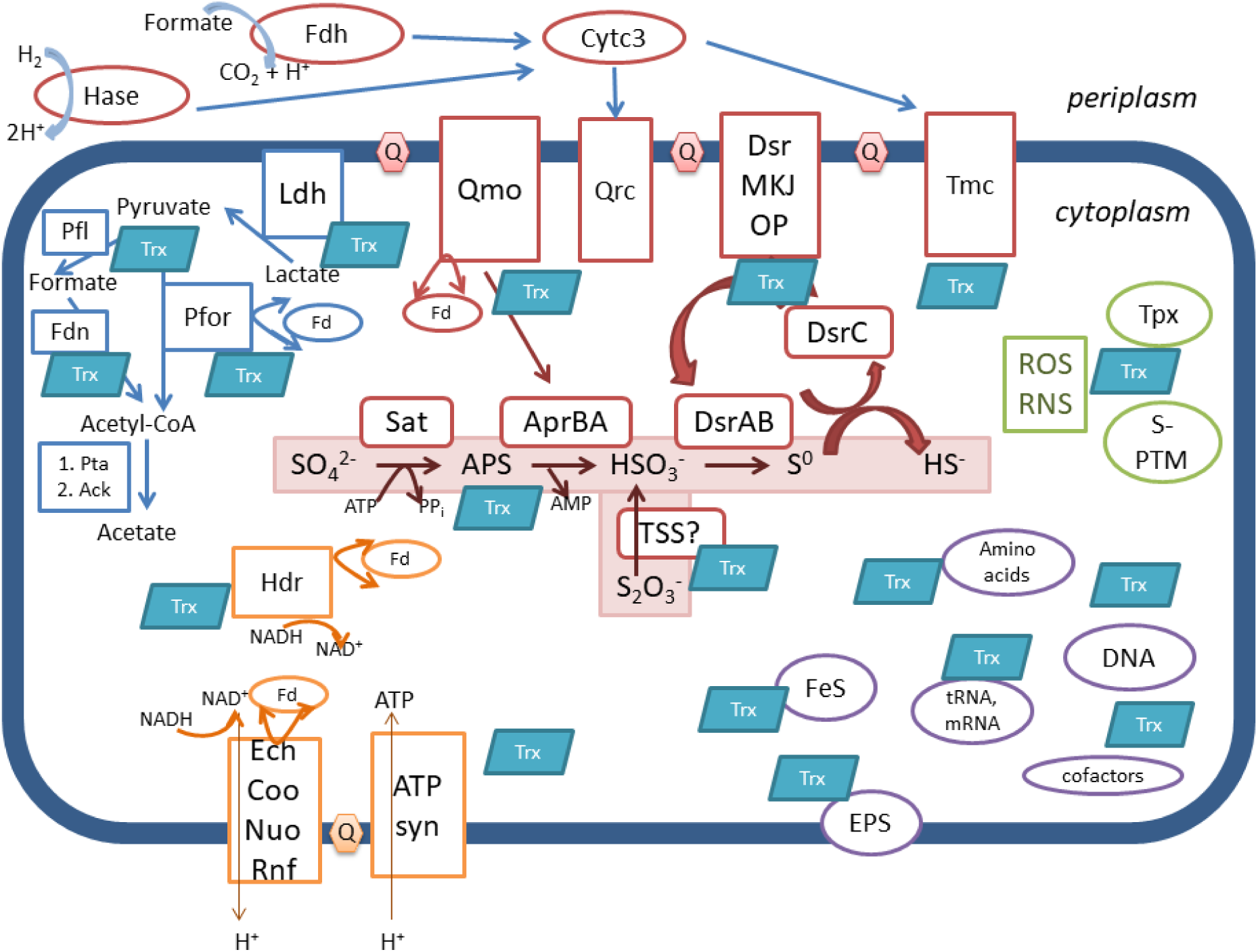
Summary model of thioredoxin involvement in SRM metabolism. Energy generating proteins are outlined in red (cytochrome involved) and orange, carbon metabolism is in blue, biosynthetic processes are in purple and oxidative stress response is in green. Hase- hydrogenase; Fdh- formate dehydrogenase; Cytc3- cytochrome c_3_; Tmc- transmembrance complex; DsrMKJOP, DsrC, DsrAB- dissimilatory sulfite reductase complex; Qrc- quinone reductase complex; Qmo- quinone monooxygenase complex; Ldh- lactate dehydrogenase; Pfor- pyruvate:ferredoxin oxidoreductase; Pfl- pyruvate formate lyase; Fdn- formate dehydrogenase; Pta- phosphate acetyltransferase; Ack- acetate kinase; Hdr- heterodisulfide reductase; Ech, Coo, Nuo, Rnf- predicted proton pumping membrane proteins; ATP syn- ATP synthase; Sat- ATP sulfurylase; AprBA- APS reductase; FeS- iron sulfur cluster; EPS- extracellular polymeric sugar; ROS/RNS- reactive oxygen/nitrogen species; Tpx- peroxiredoxin; S-PTM- thiol mediated post translational modification.

## 5. Conclusion

Taken together, our data demonstrate that SRMs lack common redox maintenance molecules and macromolecules and use a different way to maintain redox balance in their cells. SRMs have diverse thioredoxins, which are required and universal in their metabolism. Trx1 plays essential roles in transcription, translation, and cofactor biosynthesis. Trx3, an atypical thioredoxin, play roles in sulfate reduction and protein translation. To the best of our knowledge, this is the first report of substrate proteins for a thioredoxin with an atypical active site. Moreover, thioredoxins are acting in DSR, with the assistance of excreted sulfide, serving as the oxidative stress response and maintainers of redox homeostasis in SRMs. Resistance to metal stress in DvH was related to mode of respiration and presence of excreted sulfide. Sulfide may bind the toxic metals and prevent the metals from inhibiting thioredoxins or exerting its toxic effects to DvH. We also explored the novel roles of thioredoxins in SRMs such as handling ROS and RNS, involving DSR and acting as alternative biosynthesis of methionine. This could further improve the understanding of the functions of thioredoxin in SRM physiology.

## Acknowledgements

This material by ENIGMA - Ecosystems and Networks Integrated with Genes and Molecular Assemblies (http://enigma.lbl.gov), a Science Focus Area Program at Lawrence Berkeley National Laboratory is based upon work supported by the U.S. Department of Energy, Office of Science, Biological & Environmental Research Program under contract number DE-AC02-05CH11231. The running of the proteomics samples was supported by the University of Missouri Proteomics Core Grant, and we thank Proteomics Core staff, especially Brian Mooney, for helping with the sample preparation and raw data analysis. We thank Kara B. De León, Grant M. Zane, Thomas R. Juba, and Kinjal Majumder for their assistance with discussions, using their matching, genome assembly, and growth curve plotting scripts and providing mutants which served as templates for the ones generated in this study.

